# Chytrid rhizoid morphogenesis is adaptive and resembles hyphal development in ‘higher’ fungi

**DOI:** 10.1101/735381

**Authors:** Davis Laundon, Nathan Chrismas, Glen Wheeler, Michael Cunliffe

## Abstract

Fungi are major components of the Earth’s biosphere [1], sustaining many critical ecosystem processes [2, 3]. Key to fungal prominence is their characteristic cell biology, our understanding of which has been principally based on ‘higher’ dikaryan hyphal and yeast forms [4–6]. The early-diverging Chytridiomycota (chytrids) are ecologically important [2, 7, 8] and a significant component of fungal diversity [9–11], yet their cell biology remains poorly understood. Unlike dikaryan hyphae, chytrids typically attach to substrates and feed osmotrophically via anucleate rhizoids [12]. The evolution of fungal hyphae appears to have occurred from lineages exhibiting rhizoidal growth [13] and it has been hypothesised that a rhizoid-like structure was the precursor to multicellular hyphae and mycelial feeding in fungi [14]. Here we show in a unicellular chytrid, *Rhizoclosmatium globosum*, that rhizoid development has equivalent features to dikaryan hyphae and is adaptive to resource availability. Rhizoid morphogenesis exhibits analogous properties with growth in hyphal forms, including tip production, branching and decreasing fractal geometry towards the growing edge, and is controlled by β-glucan-dependent cell wall synthesis and actin polymerisation. Chytrid rhizoids from individual cells also demonstrate adaptive morphological plasticity in response to substrate availability, developing a searching phenotype when carbon starved and exhibiting spatial differentiation when interacting with particulate substrates. Our results show striking similarities between unicellular early-diverging and dikaryan fungi, providing insights into chytrid cell biology, ecological prevalence and fungal evolution. We demonstrate that the sophisticated cell biology and developmental plasticity previously considered characteristic of hyphal fungi are shared more widely across the Kingdom Fungi and therefore could be conserved from their most recent common ancestor.

## Introduction

The phylum Chytridiomycota (chytrids) diverged approximately 750 million years ago and, with the Blastocladiomycota, formed a critical evolutionary transition in the Kingdom Fungi dedicated to osmotrophy and the establishment of the chitin-containing cell wall [10]. 407-million-year-old chytrid fossils from the Devonian Rhynie Chert deposit show chytrids physically interacting with substrates via rhizoids in a comparative way to extant taxa [15]. Rhizoids play key roles in chytrid ecological function, in terms of both attachment to substrates and osmotrophic feeding [10, 12]. Yet surprisingly, given the importance of rhizoids in chytrid ecology, there remains a lack of understanding of chytrid rhizoid biology, including potential similarities with the functionally analogous hyphae in other fungi.

While both rhizoids and hyphae are polar, elongated and bifurcating structures, rhizoid feeding structures are a basal condition within the true fungi (Eumycota), and the dikaryan mycelium composed of multicellular septate hyphae is highly derived (Figure 1A and B). Hyphal cell types are observed outside of the Eumycota, such as within the Oomycota, however the origin of fungal hyphae within the Eumycota was independent [13, 16] and has not been reported in their closest relatives the Holozoans (animals, choanoflagellates and their kin). Comparative genomics has indicated that hyphae originated within the Chytridiomycota-Blastocladiomycota-Zoopagomycota nodes of the fungal tree [16], and is supported by fossil Blastocladiomycota and extant Monoblepharidomycetes having hyphae [13, 17]. However, even though rhizoids have been considered precursory to hyphae [14], comparisons between rhizoid and hyphal developmental biology have not yet been made.

**Figure 1.**
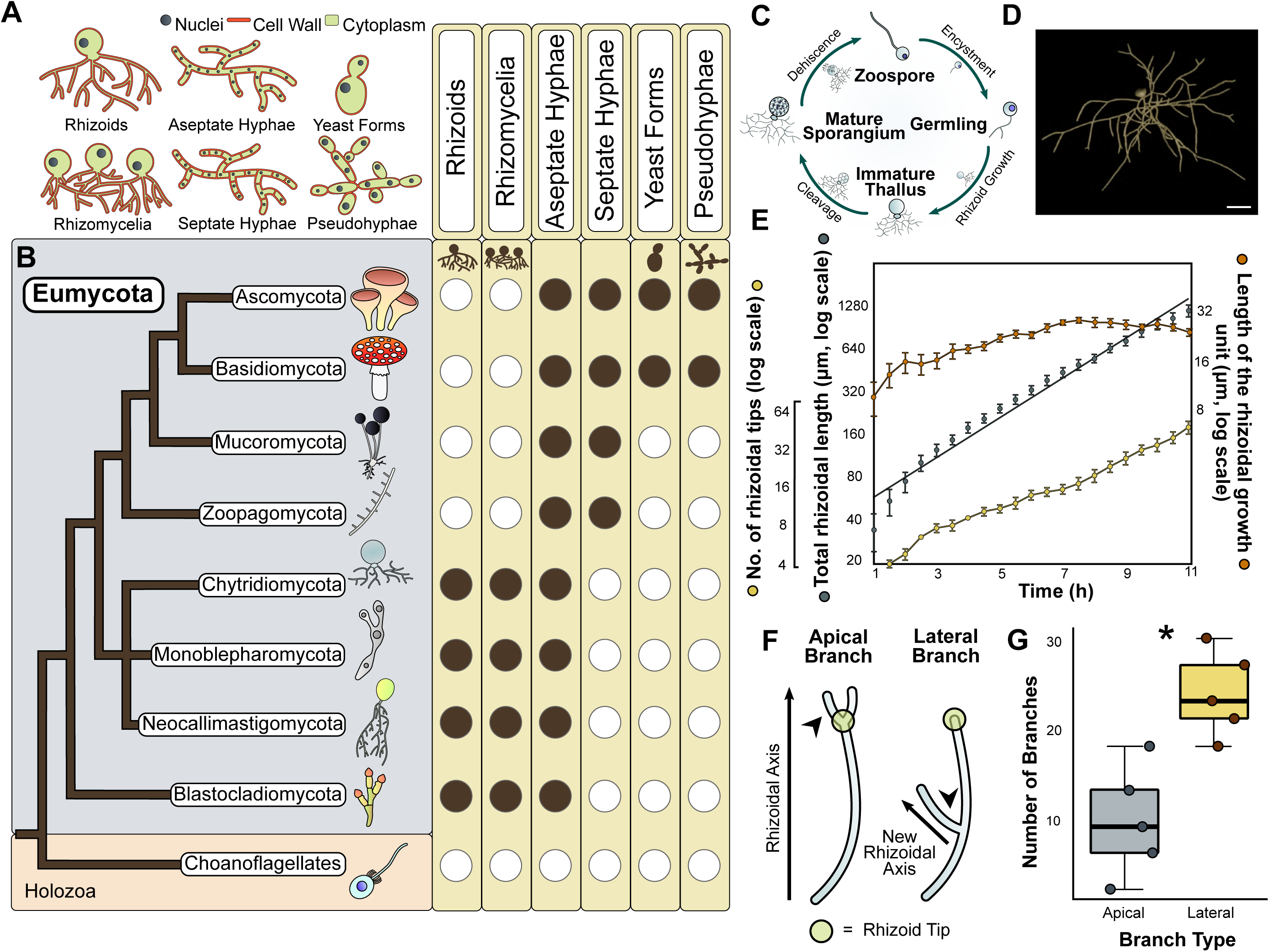
Rhizoids are the basal feeding condition within the fungal kingdom and their morphogenesis is similar to hyphal development. (A-B) Correlating the major feeding types in fungi (A) to phylogeny (B) shows rhizoids to be the basal feeding condition in the true fungi (Eumycota). Tree adapted from [11]. (C) *R. globosum* exhibits an archetypal chytrid lifecycle. (D) Chytrid rhizoids were reconstructed using the neuron tracing workflow outlined in Supplementary Figure 3. Example of a 3D reconstructed *R. globosum* rhizoid system taken from a 10 h time series. Scale bar = 20 µm. (E) Rhizoid growth trajectories for 4D confocal time series (*n* = 5, mean ± S.E.M.) of rhizoidal growth unit, total length and number of tips. (F) Apical and lateral branches occur in chytrid rhizoids. Apical branching occurs when a branch is formed at the rhizoid tip parallel to the established rhizoidal axis. Lateral branching occurs when a branch is formed distally to the rhizoidal tip, establishing a new rhizoidal axis. (G) 4D confocal imaging (*n* = 5, mean ± S.E.M.) revealed that lateral branching dominates over apical branching \**p* < 0.05.

*R. globosum* JEL800 is a monocentric eucarpic chytrid, with extensive anucleate thin rhizoids (230.51 ± 62.40 nm in width; Supplementary Figures 1 and 2) and an archetypal chytrid lifecycle (Figure 1C). With an available sequenced genome [18], easy laboratory culture and amenability to live cell imaging (this study), *R. globosum* represents a promising new model organism to investigate the cell biology of rhizoid-bearing, early-diverging fungi. To study the developing rhizoid system for morphometric analyses, we established a live cell 3D/4D confocal microscopy approach in combination with neuron tracing software to 3D reconstruct developing cells (Figure 1D; Supplementary Figures 3 and 4). From these reconstructions, we were able to generate a series of cell morphometrics adapted from neuronal biology to describe and quantify rhizoid development (Supplementary Figure 5).

## Results and Discussion

### Chytrid rhizoid morphogenesis fundamentally resembles mycelial development

During rhizoid development we observed a continuous increase in rhizoid length (110.8 ± 24.4 µm h^−1^) (*n* = 5, ± SD) and the number of rhizoid tips (4.6 ± 1.2 tips h^−1^) (Figure 1E; Supplementary Table 1; Supplementary Movies 1-5), with a continuous increase in the thallus surface area (21.1 ± 5.2 µm^2^ h^−1^), rhizoid bifurcations (4.2 ± 1.0 bifurcations h^−1^), cover area (2,235 ± 170.8 µm^2^ h^−1^) and maximum Euclidean distance (5.4 ± 0.1 µm h^−1^) (Supplementary Figure 6). The rhizoidal growth unit (RGU) (i.e. the distance between two rhizoid compartments) increased continuously during the first 6 h of the development period (i.e. cells became relatively less branched) before stabilising during the later phase of growth (Figure 1E).

The RGU patterns that we report here for a unicellular non-hyphal fungus are comparable to the hyphal growth units (HGU) recorded in multicellular hyphal fungi (Supplementary Figure 7) [19]. Trinci (1974) assessed hyphal development in three major fungal lineages (Ascomycota, Basidiomycota, Mucoromycota) and observed that the growth patterns of major morphometric traits (HGU, total length and number of tips) were similar across the studied taxa. When the data from our study are directly compared to that of Trinci (1974), we see that the hyphal growth pattern is also analogous to the rhizoids of the early-diverging unicellular Chytridiomycota (Supplementary Figure 7).

In *R. globosum*, the local rhizoid bifurcation angle remained consistent at 81.4° ± 6.3 after ∼2 h (Supplementary Figure 6), suggesting the presence of a currently unknown control mechanism regulating rhizoid branching in chytrids. During rhizoid development, lateral branching was more frequent than apical branching (Figure 1F and G), as observed in dikaryan hyphae [20]. Fractal analysis (fractal dimension = D*b*) of 24 h chytrid cells revealed that rhizoids approximate a 2D biological fractal (Mean D*b* = 1.51 ± 0.24), with rhizoids relatively more fractal at the centre of the cell (Max D*b* = 1.69-2.19) and less fractal towards the growing periphery (Min D*b* = 0.69-1.49) (Supplementary Figure 8). Similar patterns of fractal organisation are also observed in hyphae-based mycelial colonies [21]. Together these findings suggest that a form of apical dominance at the growing edge rhizoid tips may suppress apical branching to maintain rhizoid network integrity as in dikaryan hyphae [22, 23].

### Cell wall and actin dynamics govern branching in chytrid rhizoids

Given the apparent hyphal-like properties of the chytrid cell, we sought a greater understanding of the potential subcellular machinery underpinning rhizoid morphogenesis. Chemical characterisation of the *R. globosum* rhizoid showed that the chitin-containing cell wall and actin patches were located throughout the rhizoid (Figure 2A). As the cell wall and actin control hyphal morphogenesis in dikaryan fungi [4–6], they were selected as targets for chemical inhibition in the chytrid. Inhibition of cell wall β-1,3-glucan synthesis and actin proliferation with caspofungin and cytochalasin B respectively induced a concentration-dependent decrease in the RGU and the development of atypical cells with hyperbranched rhizoids (Figures 2B-D; Supplementary Table 2; Supplementary Movies 6-7). These effects in *R. globosum* are similar to disruption of normal hyphal branching reported in *Aspergillus fumigatus* (Ascomycota) in the presence of caspofungin [24], and in *Neurospora crassa* (Ascomycota) in the presence of cytochalasins [25], suggesting that β-1,3-glucan-dependent cell wall synthesis and actin dynamics also govern branching in chytrid rhizoids by comparable processes.

**Figure 2.**
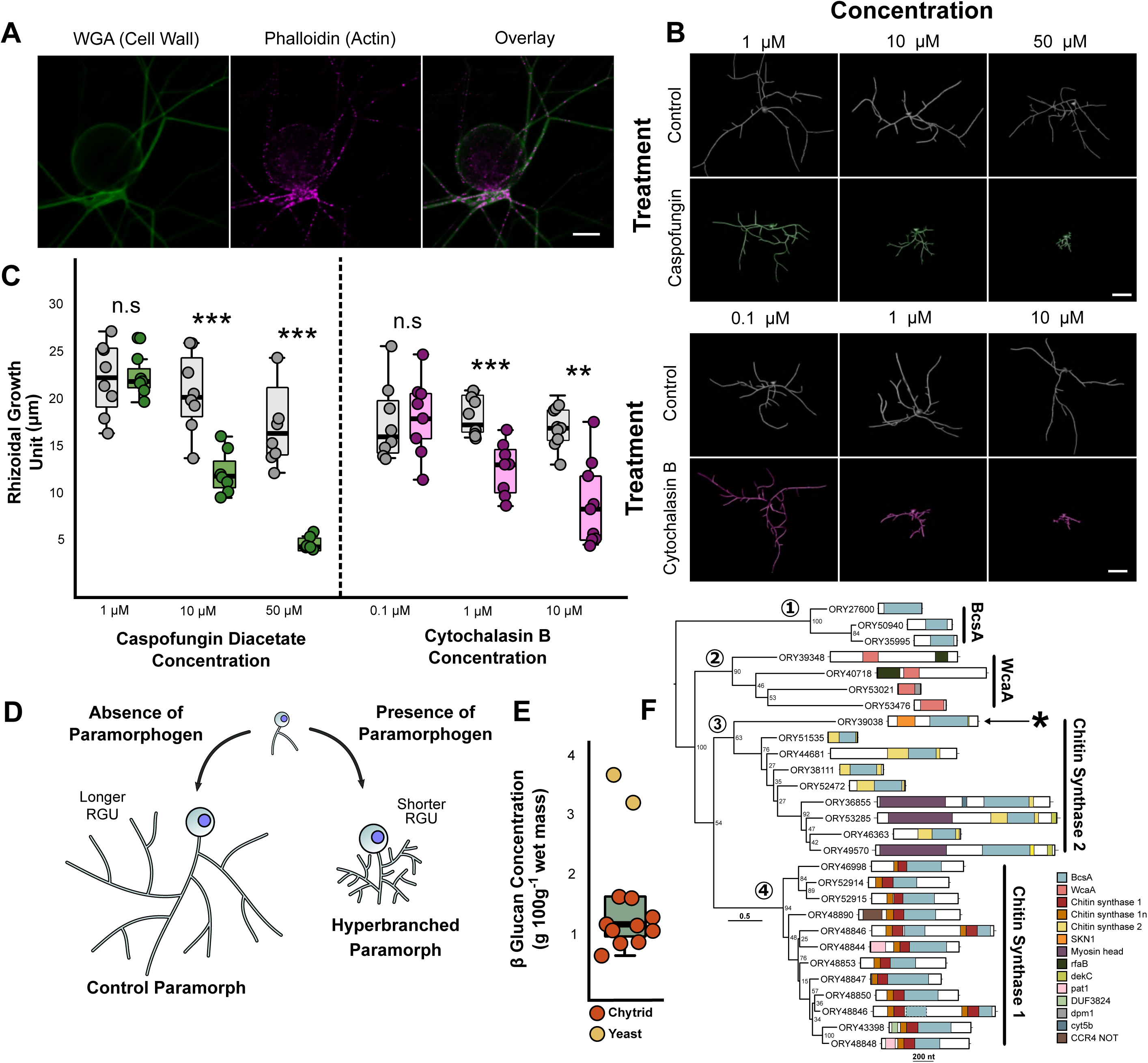
Cell wall synthesis and actin dynamics govern rhizoid branching. (A) Fluorescent labelling of cell wall and actin structures in 24 h *R. globosum* cells. The cell wall and actin patches were found throughout the rhizoid. WGA = conjugated Wheat Germ Agglutinin. Scale bar = 10 µm. (B) Representative 3D reconstructions of 7 h *R. globosum* cells following treatment with caspofungin diacetate and cytochalasin B at stated concentrations to inhibit cell wall and actin filament biosynthesis respectively, relative to solvent only controls. Scale bar = 20 µm (C) Application of caspofungin diacetate and cytochalasin B resulted in a concentration-dependent decrease in the rhizoidal growth unit, resulting in atypical hyperbranched rhizoids (*n* ∼9, mean ± S.E.M.). n.s *p* > 0.05 (not significant), \**p* < 0.05, \*\**p* < 0.01, \*\*\**p* < 0.001. This differential growth is diagrammatically summarised in (D). (E) β-glucan concentration of *R. globosum* (*n* = 10) relative to a baker’s yeast control (*n* = 2). (F) Maximum likelihood phylogeny of GT2 domains (BcsA and WcaA domains) within the *R. globosum* genome (midpoint rooting). Full architecture of each protein is shown. Asterisk indicates the putative glucan synthesis protein ORY39038 containing a putative SKN1 domain.

*In silico* studies of fungal genomes have proposed that the Chytridiomycota (represented by *Batrachochytrium dendrobatidis*) lack β-1,3-glucan synthase FSK1 gene homologs [26–28], which is the target for caspofungin. Despite the absence of FKS1 homologues in chytrid genomes, quantification of glucans in *R. globosum* showed that they are present (Figure 2E), with 58.3 ± 7.6 % β-glucans and 41.6 ± 7.6 % α-glucans of total glucans.

To identify putative β-glucan synthesis genes, we surveyed the *R. globosum* JEL800 genome and focused on glycosyltransferase family 2 (GT2) encoding genes, which include typical glucan synthases in fungi. A total of 28 GT2 domains were found within 27 genes (Figure 2F). Of these genes, 20 contained putative chitin synthase domains and many contained additional domains involved in transcriptional regulation. Nine encode chitin synthase 2 family proteins and 11 encode chitin synthase 1 family proteins (with two GT2 domains in ORY48846). No obvious genes for β-1,3-glucan or β-1,6-glucan synthases were found within the genome, consistent with previous *B. dendrobatidis* studies [27, 28].

However, the chitin synthase 2 gene ORY39038 included a putative SKN1 domain (Figure 2F), which has been implicated in β-1,6-glucan synthesis in the ascomycete yeasts *Saccharomyces cerevisiae* [29] and *Candida albicans* [30]. These results indicate a yet uncharacterised β-glucan-dependent cell wall production process in chytrids (also targeted by caspofungin) that is not currently apparent using gene/genome level assessment and warrants further study.

### Chytrid rhizoids undergo adaptive development in response to carbon starvation

To examine whether chytrids are capable of modifying rhizoid development in response to changes in resource availability, we exposed *R. globosum* to carbon starvation (i.e. development in the absence of exogenous carbon). When provided with 10 mM *N*-acetyl-D-glucosamine (NAG) as an exogenous carbon source, the entire life cycle from zoospore to sporulation was completed (Supplementary Movie 8). Carbon-starved cells did not produce zoospores and cell growth stopped after 14-16 h (Supplementary Movie 9). However, using only endogenous carbon (i.e. zoospore storage lipids) carbon starved cells underwent substantially differential rhizoid development compared to cells from the exogenous carbon replete conditions that we interpret to be an adaptive searching phenotype (Figure 3A and B; Supplementary Table 4; Supplementary Movie 10). Under carbon starvation, *R. globosum* cells invested less in thallus growth than in carbon replete conditions, with the development of longer rhizoids with a greater maximum Euclidean distance (Figure 3C). Carbon starved cells were also less branched, had wider bifurcation angles and subsequently covered a larger surface area. These morphological changes in response to exogenous carbon starvation (summarised in Figure 3B) suggest that individual chytrid cells are capable of controlled reallocation of resources away from reproduction (i.e. the production of the zoosporangium) and towards an extended modified rhizoidal structure indicative of a resource searching phenotype. Exogenous carbon starvation has also been shown to be associated with a decrease in branching in the multicellular dikaryan fungus *Aspergillus oryzae* (Ascomycota) [31]. Branching zones in dikaryan mycelia are known to improve colonisation of trophic substrates and feeding, while more linear ‘exploring’ zones search for new resources [32].

**Figure 3.**
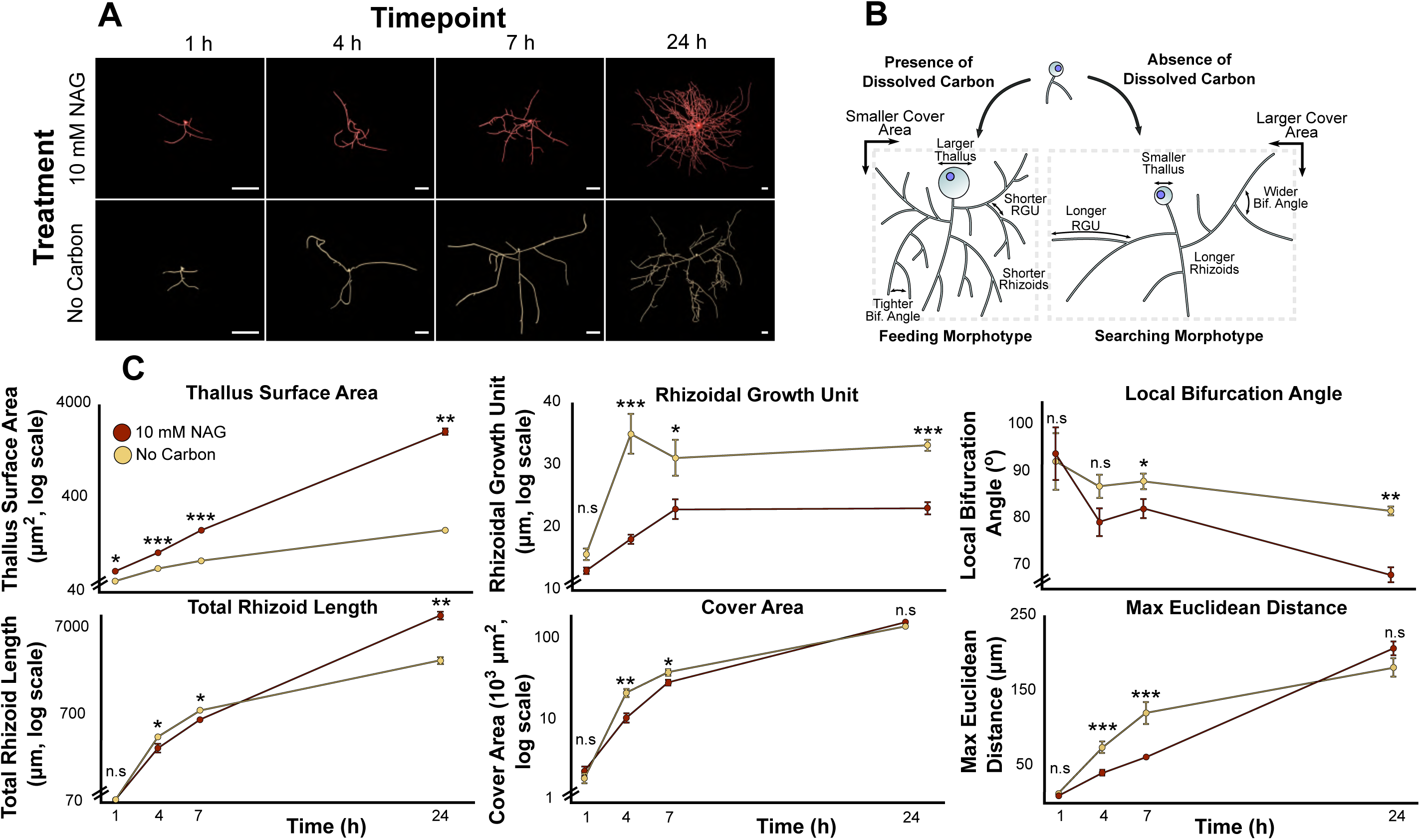
Chytrids are capable of adaptive rhizoid development under carbon starvation. (A) Representative 3D reconstructions of *R. globosum* cells grown under carbon replete or carbon deplete conditions at different timepoints. Scale bar = 20 µm. When exposed to carbon starvation, chytrids are capable of differential adaptive growth to produce a searching phenotype. This differential growth is summarised in (B). (C) Differential growth trajectories of major morphometric traits between *R. globosum* cells (*n* ∼9, mean ± S.E.M.) grown under carbon replete and carbon deplete conditions over time. n.s *p* > 0.05 (not significant), \**p* < 0.05, \*\**p* < 0.01, \*\*\**p* < 0.001

### Chytrids exhibit spatially differentiated rhizoids in response to patchy environments

In the natural environment, chytrids inhabit structurally complex niches made up of heterologous substrates, such as algal cells [33], amphibian epidermises [34] and recalcitrant particulate organic carbon [35]. *R. globosum* is a freshwater saprotrophic chytrid that is typically associated with chitin-rich insect exuviae [36]. We therefore quantified rhizoid growth of single cells growing on chitin microbeads as an experimental particulate substrate (Figure 4A and B; Supplementary Movie 11). Initially, rhizoids grew along the outer surface of the bead and were probably used primarily for anchorage to the substrate. Scanning electron microscopy (SEM) showed that the rhizoids growing externally on the chitin particle formed grooves on the bead parallel to the rhizoid axis (Supplementary Figure 1F and G), suggesting extracellular enzymatic chitin degradation by the rhizoid on the outer surface. Penetration of the bead occurred during the later stages of particle colonisation (Figure 4A; Supplementary Movie 12). Branching inside the bead emanated from ‘pioneer’ rhizoids that penetrated into the particle (Figure 4C).

**Figure 4.**
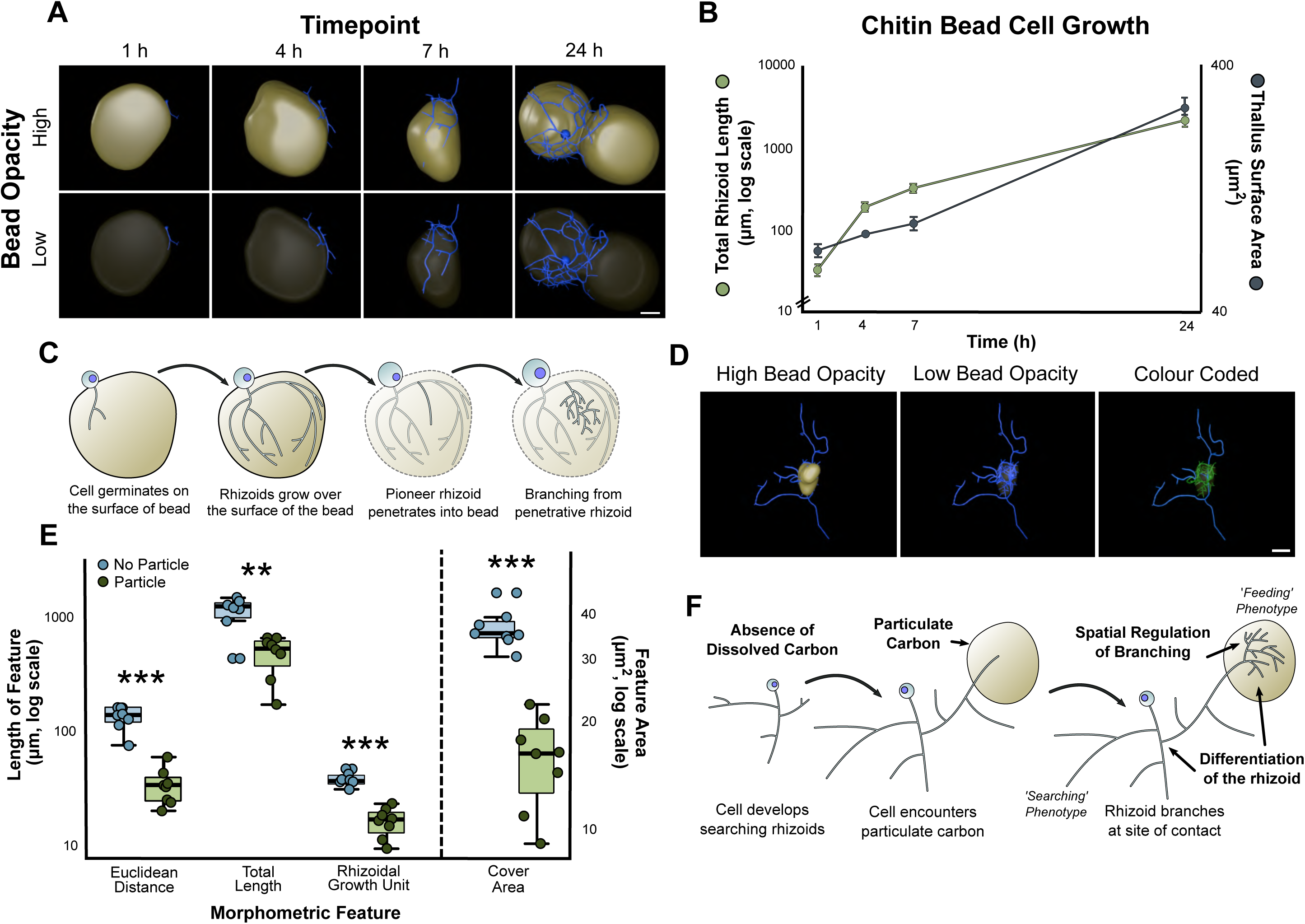
Rhizoids associated with heterogenous particulate carbon exhibit spatial differentiation. (A) Representative 3D reconstructions of *R. globosum* cells (blue) growing on chitin beads (beige) at different timepoints. Scale bar = 20 µm. (B) Growth trajectories for total rhizoid length and thallus surface area for *R. globosum* cells growing on chitin beads (*n*∼9, mean ± S.E.M.). (C) Diagrammatic summary of *R. globosum* rhizoid development on chitin beads. (D) Representative 3D reconstruction of a 24 h searching *R. globosum* cell (blue) that has encountered a chitin bead (beige). The colour coded panel shows parts of the rhizoid system in contact (green) and not in contact (blue) with the bead. Scale bar = 20 µm. (E) Comparison of rhizoids in contact or not in contact with the chitin bead (*n* = 8, mean ± S.E.M.). (F) Diagrammatic summary of spatial differentiation in a starved, searching rhizoid that has encountered a particulate carbon patch.

Given the previous results of the searching rhizoid development in response to carbon starvation, we created a patchy resource environment using the chitin microbeads randomly distributed around individual developing cells in otherwise carbon-free media to investigate how encountering a carbon source affected rhizoid morphology (Figure 4D; Supplementary Movies 13-15). Particle-associated rhizoids were shorter than rhizoids not in particle contact, were more branched (i.e. lower RGU), had a shorter maximum Euclidean distance and covered a smaller area (Figure 4E). These simultaneous feeding and searching modifications in individual cells linked to particle-associated and non-associated rhizoids respectively are similar to the rhizoid morphometrics of the cells grown under carbon replete and carbon deplete conditions previously discussed (Figure 4F and Figure 3B). The simultaneous display of both rhizoid types in the same cell suggests a controlled spatial regulation of branching and differentiation of labour within the individual anucleate rhizoidal network. Functional division of labour is seen in multicellular mycelia fungi [32, 37], including developing specialised branching structures for increased surface area and nutrient uptake as in the plant symbiont mycorrhiza (Glomeromycota) [38]. Our observation of similar complex development in a unicellular chytrid suggests that multicellularity is not a prerequisite for adaptive spatial differentiation in fungi.

### Conclusions

Appreciation for the ecological significance of chytrids as saprotrophs, parasites and pathogens is greatly expanding. For example, chytrids are well-established plankton parasites [8], responsible for the global-scale amphibian pandemic [7] and have recently emerged as important components of the marine mycobiome [2]. The improved understanding of chytrid rhizoid biology related to substrate attachment and feeding we present here opens the door to a greater insight into the functional ecology of chytrids and their ecological potency. From an evolutionary perspective, the early-diverging fungi are a critical component of the eukaryotic tree of life [9, 39], including an origin of multicellularity and the establishment of the archetypal fungal hyphal form, which is responsible in part, for the subsequent colonisation of land by fungi, diversity expansion and interaction with plants [10]. Our cell biology focused approach advances this developing paradigm by showing that a representative monocentric, rhizoid-bearing (i.e. non-hyphal) chytrid displays hyphal-like morphogenesis, with evidence that the cell structuring mechanisms underpinning chytrid rhizoid development are equivalent to reciprocal mechanisms in dikaryan fungi. Perhaps our key discovery is that the anucleate chytrid rhizoid shows considerable developmental plasticity. *R. globosum* is able to control rhizoid morphogenesis to produce a searching form in response to carbon starvation and, from an individual cell, is capable of spatial differentiation in adaptation to patchy substrate availability indicating functional division of labour. The potential for convergent evolution aside, we conclude by parsimony from the presence of analogous complex cell developmental features in an extant representative chytrid and dikaryan fungi that adaptive rhizoids, or rhizoid-like structures, are precursory to hyphae, and are a shared feature of their most recent common ancestor.

## Methods

### Culture and maintenance

For routine maintenance, *Rhizoclosmatium globosum* JEL800 was grown on PmTG agar [40]. Agar plugs were excised from established cultures using a sterile scalpel, inverted onto new agar plates and incubated at 22 °C in the dark for 48 h. Developed zoosporangia were sporulated by covering each plug with 100 µl dH2O and incubating at room temperature for 30 min. The released zoospores were distributed across the agar surface by tilting, dried for 10 min in a laminar flow hood and incubated as above. To harvest zoospores for experiments, plates were flooded with 1 ml dH2O and the zoospore suspension passed through a 10 μm cell strainer (pluriSelect) to remove mature thalli. Zoospore density was quantified using a Sedgewick Raft Counter (Pyser SCGI) and a Leica DM1000 (10 x objective) with cells fixed in 2% formaldehyde at a dilution of 1:1,000. Zoospores were diluted to a working density of 6.6 × 10^3^ ml^−1^ for all experiments. Because PmTG is a complex medium, all experiments detailed below were conducted in Bold’s Basal Medium (BBM) supplemented with 1.89 mM ammonium sulfate and 500 µl.l^−1^ F/2 vitamin solution [41].

### General cell imaging

To visualise the rhizoids, cell plasma membranes were labelled with 8.18 μM FM® 1-43 and imaged using a Zeiss LSM 510 Meta confocal laser scanning microscope (CLSM) (Carl Zeiss) under a 40 x oil-immersion objective lens, with excitation by a 488 nm Ar laser and emission at 500-530 nm. Z-stacks were acquired at 1 µm intervals. For Scanning Electron Microscopy (SEM) of rhizoids growing along a 2D surface, culture dishes were lined with EtOH-sterilised Aclar® disks and filled with 3 ml of BBM with 10 mM NAG, before inoculation with zoospores and incubation for 24 h at 22 °C. For SEM of cells growing on chitin beads, dishes were prepared as described below and were also inoculated and incubated for 24 h. Following incubation, cells were fixed overnight in 2.5% glutaraldehyde and then rinsed twice in 0.1 M cacodylate buffer (pH 7.2). Fixed samples were dehydrated in a graded alcohol series (30%, 50%, 70%, 90%, 100%) with a 15 min incubation period between each step. Cells were then dried in a Critical Point Drier (K850, Quorum) and attached to SEM sample stubs using carbon infiltrated tabs prior to Cr sputter-coating using a sputter coating unit (Q150T, Quorum). Samples were imaged with a Field Emission Gun Scanning Electron Microscope (JSM-7001F, JEOL) operating at 10 kV. For Transmission Electron Microscopy (TEM), 24 h cells grown in suspension were fixed as previously described. The samples were secondarily fixed with osmium tetroxide (1%, in buffer pH 7.2, 0.1M) for 1 h, rinsed, and alcohol dehydrated as above. The alcohol was replaced with agar low viscosity resin through a graded resin series (30%, 50%, 70%, 100%, 100%) with 12 h intervals between each step. Samples were transferred to beem capsules and placed in an embedding oven at 60 °C overnight to enable resin polymerisation. The resulting blocks were sectioned at 50 nm intervals with an ultramicrotome (Ultracut E, Leica) using a diatome diamond knife. The sections were stained using a saturated solution of uranyl acetate (for 15 min) and Reynold’s lead citrate (15 min) before being examined using a transmission electron microscope (JEM-1400, JEOL).

### 4D rhizoid development

Glass bottom dishes (*n* = 5) containing 3 ml BBM with 10 mM NAG as the available carbon source were inoculated with 500 µl zoospore suspension. Zoospores settled for 1h prior to imaging before z-stacks to 50 µm depth were acquired at 30 min time intervals for 10 h at 22 °C. Throughout the imaging duration, an optically clear film permitting gas exchange covered the dish. Branching was counted manually from maximum intensity projected z-stacks. To quantify rhizoid fractal dimensions, cells were grown on glass bottom dishes for 24 h. Due to the large size of the 24 h cells, z-stacks were stitched together in Fiji [42] from four individual stacks. Stitched stacks (*n* = 5) were converted to maximum intensity projections, processed into binary masks by default thresholding and denoised. Local Connected Fractal Dimension (LCFD) analysis was conducted using default parameters on binary masks with the Fiji plugin FracLac [43].

### Rhizoid tracing and reconstruction

Z-stacks of rhizoids were imported into the neuron reconstruction software NeuronStudio [44, 45] and adjusted for brightness and contrast. Rhizoids were semi-automatically traced with the ‘Build Neurite’ function using the basal point of the sporangium as the rhizoidal origin. Tracing used fixed intensity thresholds input optimally for each image and rhizoids were manually curated and corrected by removing tracing artefacts (e.g. correcting for loop-splitting). Cells were discarded during quality control if the tracing was substandard, accounting for the occasional variation in sample size. Cells grown for 24 h in BBM 10 mM NAG or on chitin beads were too dense to be manually curated and therefore were automatically traced using dynamic thresholding with a minimum neurite length of 2 µm, although due to their high-density tracings should be considered imperfect. For 4D image stacks, the rhizoid was reconstructed in 3D at each 30 min interval. For particle associated and non-associated rhizoids, traced rhizoid systems from individual cells were manually split into their respective categories.

Rhizoids were exported as SWC file extensions [46] and morphometrically quantified using the btmorph2 library [47] run with Python 3.6.5 implemented in Jupyter Notebook 4.4.0. Reconstructed rhizoids were visualised by converting the SWC files first to VTK files using the swc2vtk Python script (Daisuke Miyamoto: github.com/ DaisukeMiyamoto /swc2vtk/) and then to OBJ files using the ‘Extract Surface’ filter in ParaView [48]. OBJ files were then imported into Blender (2.79), smoothed using automatic default parameters and rendered for display. OBJ meshes were used for final display only and not analysis. To visualise chitin beads, z-stacks were imported into the Fiji plugin TrakEM2 [49]. Chitin beads were manually segmented, and 3D reconstructed by automatically merging traced features along the z-axis. Meshes were then preliminarily smoothed in TrakEM2 and exported as OBJ files into Blender for visualisation.

### Chemical characterisation of the rhizoid

To label the cell wall and F-actin throughout the rhizoid system, cells were grown for 24 h in 3 ml BBM with 10 mM NAG on glass bottom dishes. The culture medium was aspirated from the cells, which were then washed three times in 500 µl 1 x PBS (phosphate buffered saline). Cells were subsequently fixed for 1 h in 4% formaldehyde in 1 x PBS and then washed three times in 1 x PBS and once in PEM (100 mM PIPES (piperazine-N,N′-bis(2-ethanesulfonic acid)) buffer at pH 6.9, 1 mM EGTA (ethylene glycol tetraacetic acid), and 0.1 mM MgSO_4_). Fixed cells were stained with 1:50 rhodamine phalloidin in PEM for 30 min, washed three times in PEM, and finally stained with 5 µg/ml Texas Red-conjugated wheat germ agglutinin (WGA) in PEM for 30 min. Stained cells were further washed three times in PEM and mounted under a glass coverslip with one drop of ProLong™ Gold Antifade Mountant (ThermoFisher). Cells were imaged using the same CLSM as described above with a 63 x oil immersion objective lens. F-Actin was imaged by excitation with a 543 nm HeNe laser and emission at 535-590 nm, and the cell wall by excitation with a 633 nm HeNe laser and emission at 650-710 nm. No dye controls were run for each excitation/emission channel.

### Chemical inhibition of rhizoid growth

Autoclaved glass coverslips (VWR) were placed in a culture dish and submerged in 3 ml BBM with 10 mM NAG. Following 1 h of incubation to allow normal zoospore settlement and germination, 1 ml of growth medium was removed from the dish and 1 ml of poison-containing media was introduced. Caspofungin diacetate (working concentration 1-50 µM) was used to inhibit cell wall β-glucan synthesis and cytochalasin B (working concentration 0.1-10 µM) was used to inhibit actin filament formation. Cells were further incubated for 6 h, which was found to be sufficient to observe phenotypic variation before being removed from the incubator and held at 4 °C prior to imaging. Coverslips were removed from the dishes using EtOH-cleaned forceps and placed cell-side down into a glass bottom dish containing 100 µl of membrane dye. 3D, as opposed to 4D imaging, was chosen to allow more replication for statistical analysis. Three plates were imaged in triplicate (*n* = 9) for each poison treatment and for solvent-only (i.e. no poison) controls.

### β-glucan quantification

*R. globosum* was grown to 250 ml in BBM with 10 mM NAG (*n* = 5) for 7 d before harvesting by centrifugation at 4,700 rpm for 10 min in 50 ml aliquots and washed in 50 ml MilliQ H_2_O. The cell pellet from each flask was processed for β-glucans in duplicate using a commercial β-Glucan assay (Yeast & Mushroom) (K-YBGL, Megazyme) following the manufacturer’s protocol. A sample of shop-bought baker’s yeast was used as a control. Glucans were quantified spectrophotometrically using a CLARIOstar® Plus microplate reader (BMG Labtech).

### Identification of putative glucan synthases genes

All glycosyl transferase group 2 (GT2) domain-containing proteins within the *R. globosum* genome were identified using the JGI MycoCosm online portal. GT2 functional domains were identified using DELTA-BLAST [50] and aligned with MAFFT [51]. Maximum Likelihood phylogenies were calculated with RAxML [52] using the BLOSUM62 matrix and 100 bootstrap replicates and viewed in FigTree (Andrew Rambaut: github.com/rambaut/figtree/). Overall protein architecture was displayed using genoplotR [53].

### Carbon starvation and growth on chitin beads

To quantify differential rhizoidal growth under carbon replete and carbon deplete conditions, coverslips were placed in a culture dish and submerged in 3 ml growth medium (either carbon-free BBM or BBM with 10 mM NAG). Dishes were then inoculated with zoospores and incubated for either 1, 4, 7 or 24 h, with the 24 h cell z-stacks stitched as described in the fractal analysis. Three plates were also imaged in triplicate for each treatment at each time point (*n* = 9). For both sets of experiments, cells were imaged as per the chemical inhibition experiments above.

Chitin beads (New England Biolabs) were washed three times in carbon-free BBM using a magnetic Eppendorf rack and suspended in carbon-free BBM at a working concentration of 1:1,000 stock concentration. Glass bottom dishes containing 3 ml of the diluted beads were inoculated with zoospores and incubated for either 1, 4, 7 or 24 h prior to imaging. For imaging, the culture medium was aspirated off and beads were submerged in 100 µl FM® 1-43. Three plates were imaged in triplicate for each time point (*n* = 9). To understand rhizoid development in a starved cell that had encountered a chitin bead, we imaged cells that contacted a chitin bead following development along the glass bottom of the dish.

### Statistical Analysis

Rhizoid width was measured from TEM images (*n* = 25). The comparison between apical and lateral branching was conducted using a Wilcoxon Rank Sum test. Univariate differences in rhizoid morphometrics between experimental treatments were evaluated using Welch’s t-tests unless stated otherwise. Shapiro-Wilk and Levene’s tests were used to assess normality and homogeneity of variance respectively. If these assumptions could not be met, then Wilcoxon Rank Sum was used as a nonparametric alternative. Univariate morphometric differences between particle-associated and non-associated rhizoids were evaluated using a paired t-test. All data were analysed in RStudio v1.1.456. [54]

## Data availability

All data that support the findings of this study are freely available via the corresponding author.

## Acknowledgements

The authors would like to thank Glenn Harper, Alex Strachan and the team at the Plymouth Electron Microscopy Centre (PEMC) for their assistance. We are indebted to Joyce Longcore (University of Maine) for providing *R. globosum* JEL800 from her chytrid culture collection (now curated by the Collection of Zoosporic Eufungi at the University of Michigan).

## Funding

D.L. is supported by an EnvEast Doctoral Training Partnership (DTP) PhD studentship funded from the UK Natural Environment Research Council (NERC). M.C. is supported by the European Research Council (ERC) (MYCO-CARB project; ERC grant agreement number 772584). N.C. is supported by NERC (Marine-DNA project; NERC grant number NE/N006151/1). G.W. is supported by an MBA Senior Research Fellowship.

## Author Contributions

D.L. and M.C. conceived the study. D.L. conducted the laboratory work and data analysis.

N.C. analysed the *R. globosum* JEL800 genome. G.W. provided support with microscopy.

M.C. secured the funding. D.L. and M.C. critically assessed and interpreted the findings. D.L and M.C. wrote the manuscript, with the help of N.C. and G.W.

## Competing Interests

The authors declare no competing interests

---

**Supplementary Figure 1.**
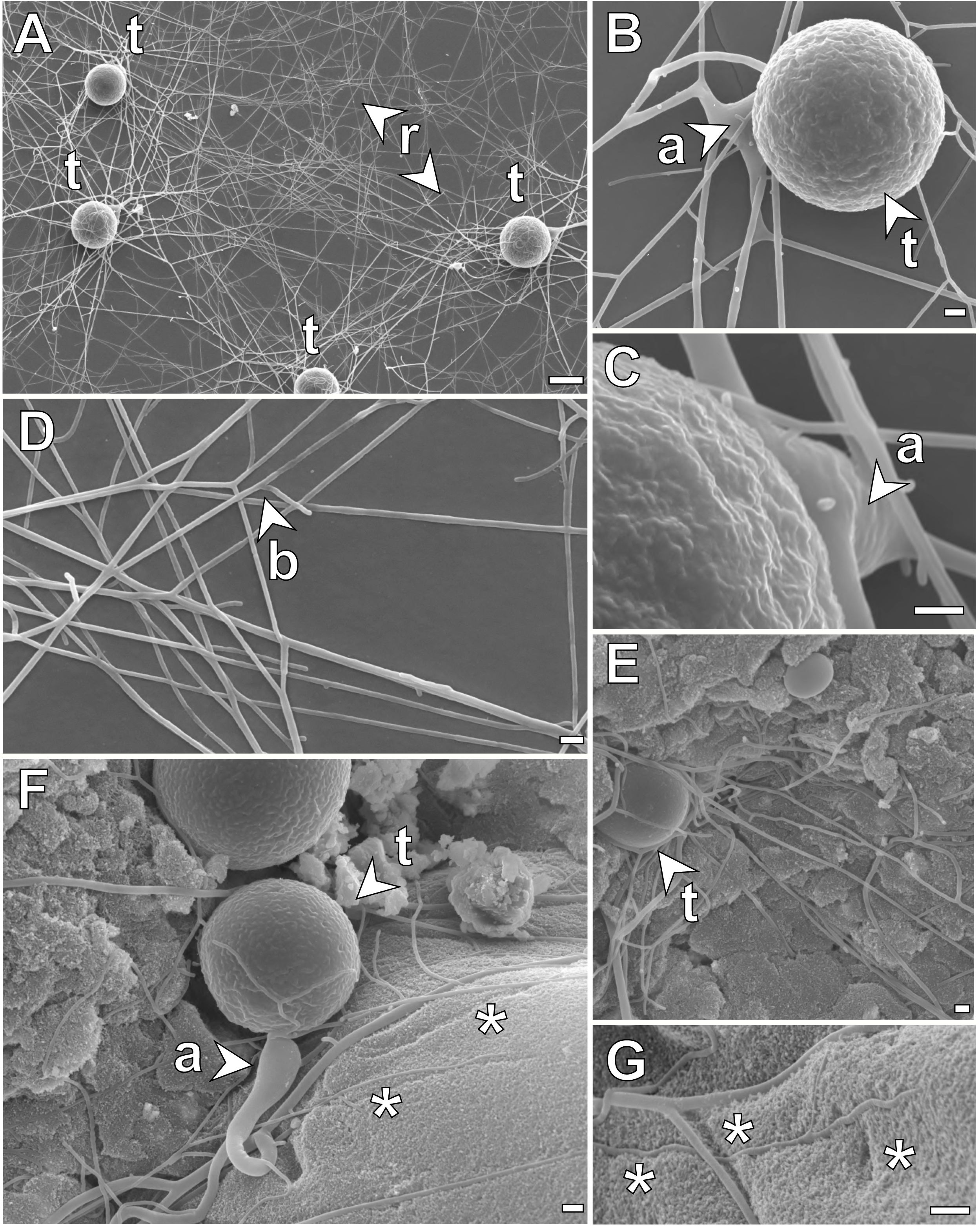
Scanning Electron Microscopy (SEM) images of *R. globosum* rhizoids. (A-D) *R. globosum* cells grown on a 2D, inert surface (Aclar®) in NAG supplemented media. (A) Shown are multiple thalli anchored to the surface by threadlike rhizoids. (B) The spherical thallus of *R. globosum* is connected to the rhizoid system via an apophysis (subsporangial swelling). (C) High-magnification image of the apophysis. (D) Rhizoids are branched and bifurcating structures that frequently overlap. The fusion of rhizoids (anastomoses) was never observed from SEM images. (E-G) Chytrid cells growing on chitin beads. (F-G) External rhizoids growing along the surface of the particle formed superficial lacerations (indicated by asterisks). a, apophysis; b, bifurcation; t, thallus. Scale bar (A,E) = 10 µm. Scale bar (B-D, F-G) = 1 µm.

**Supplementary Figure 2.**
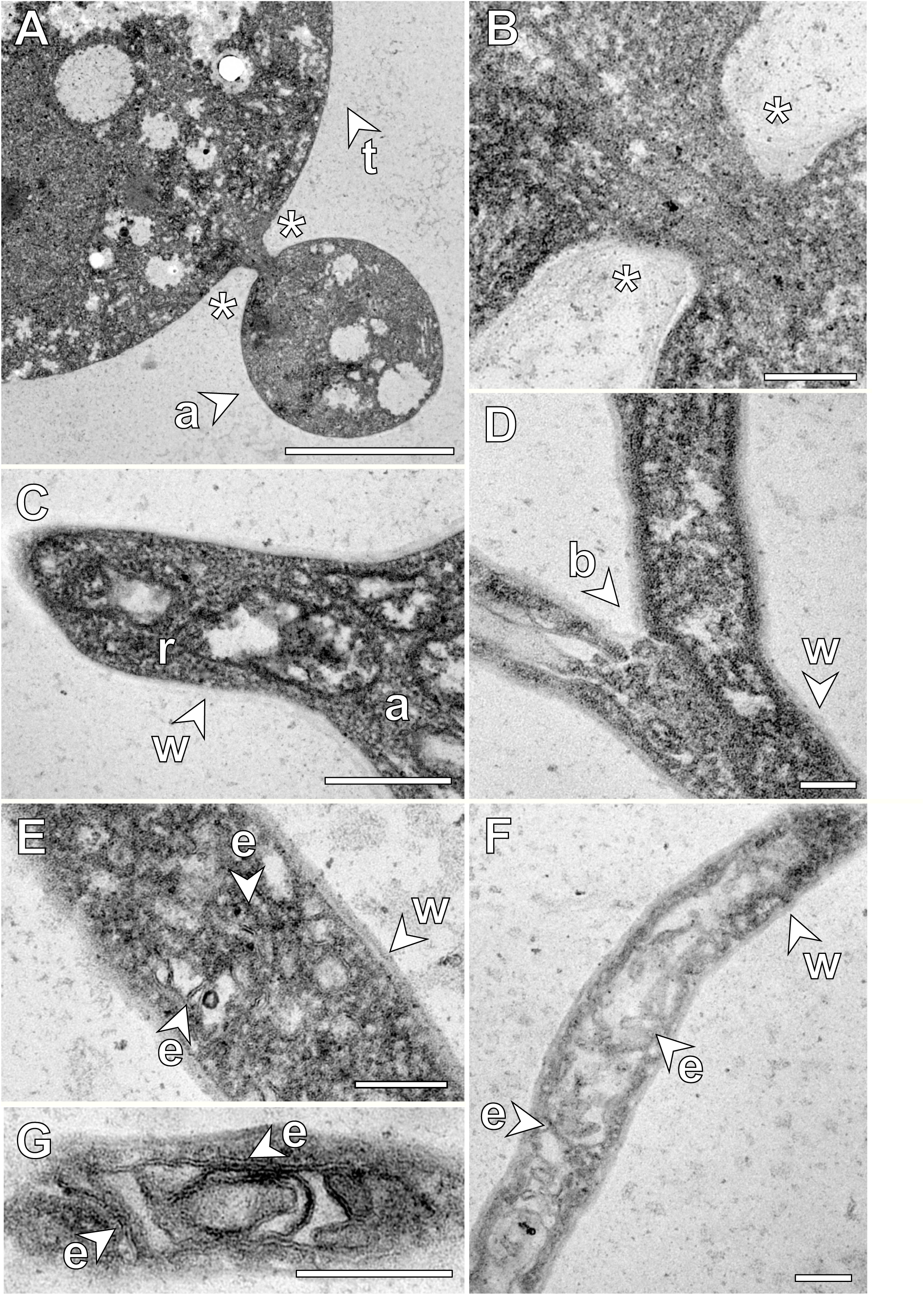
Transmission Electron Microscopy (TEM) images of *R. globosum* rhizoids. (A-C) TEM images of the apophysis. The apophysis is not septated from the thallus and the two are connected by continuous cytoplasm (A-B), as are the apophysis and the rhizoid (C). (D-F) TEM images of the apophysis. The rhizoid is always enveloped by a cell wall and no structure was observed to demarcate rhizoid branches at bifurcation nodes (D). Although no formal subcellular organelles could be identified within the rhizoid, a dense and complex endomembrane system permeated the entire system (E-F). This suggested that the rhizoid is a dynamic organelle governed by high levels of trafficking and endomembrane reorganisation. a, apohpysis; b, bifurcations; e, endomembrane; r, rhizoid; w, cell wall. Asterisks mark the connection between the apophysis and the thallus. Scale bar (A) = 2 µm. Scale bar (B-F) = 200 nm.

**Supplementary Figure 3.**
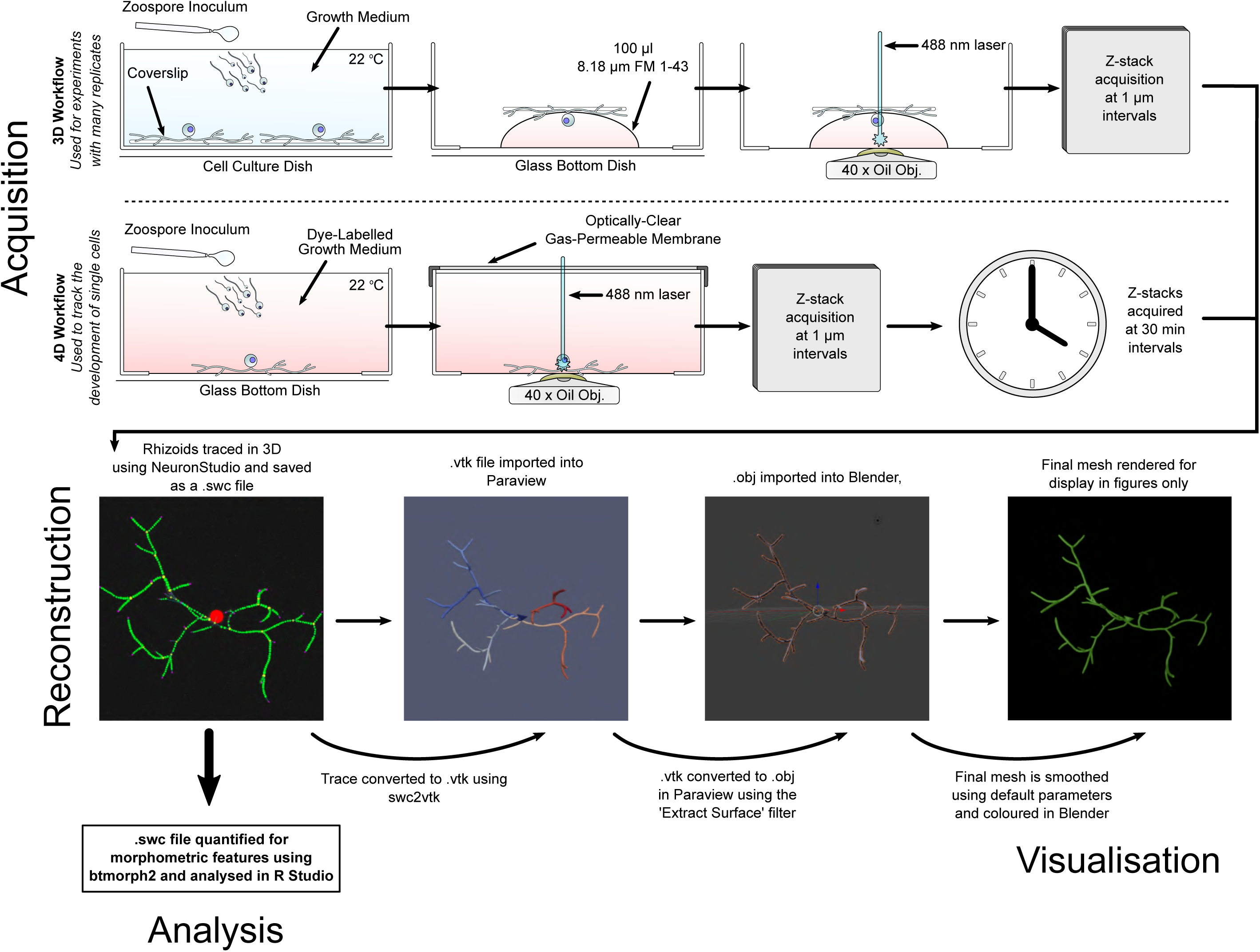
Neuron tracing was used to reconstruct and quantify chytrid rhizoid development. Flow-diagram protocol for the acquisition, reconstruction, analysis and visualisation of *R. globosum* rhizoids based on neuron tracing.

**Supplementary Figure 4.**
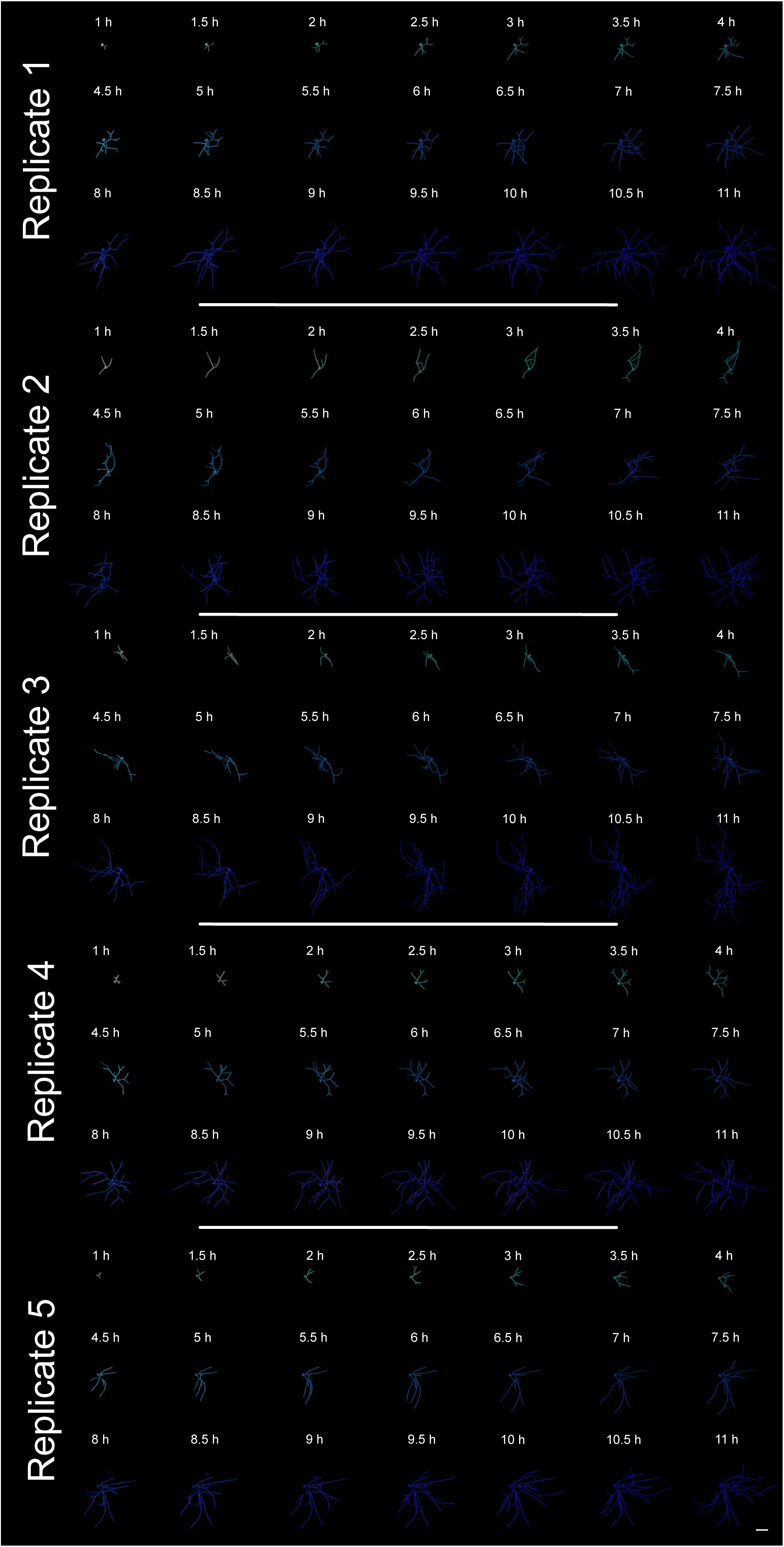
3D reconstructions of developing *R. globosum* rhizoids. Total series of 3D reconstructed *R. globosum* rhizoids taken from 4D development experiments. Scale bar = 20 µm.

**Supplementary Figure 5.**
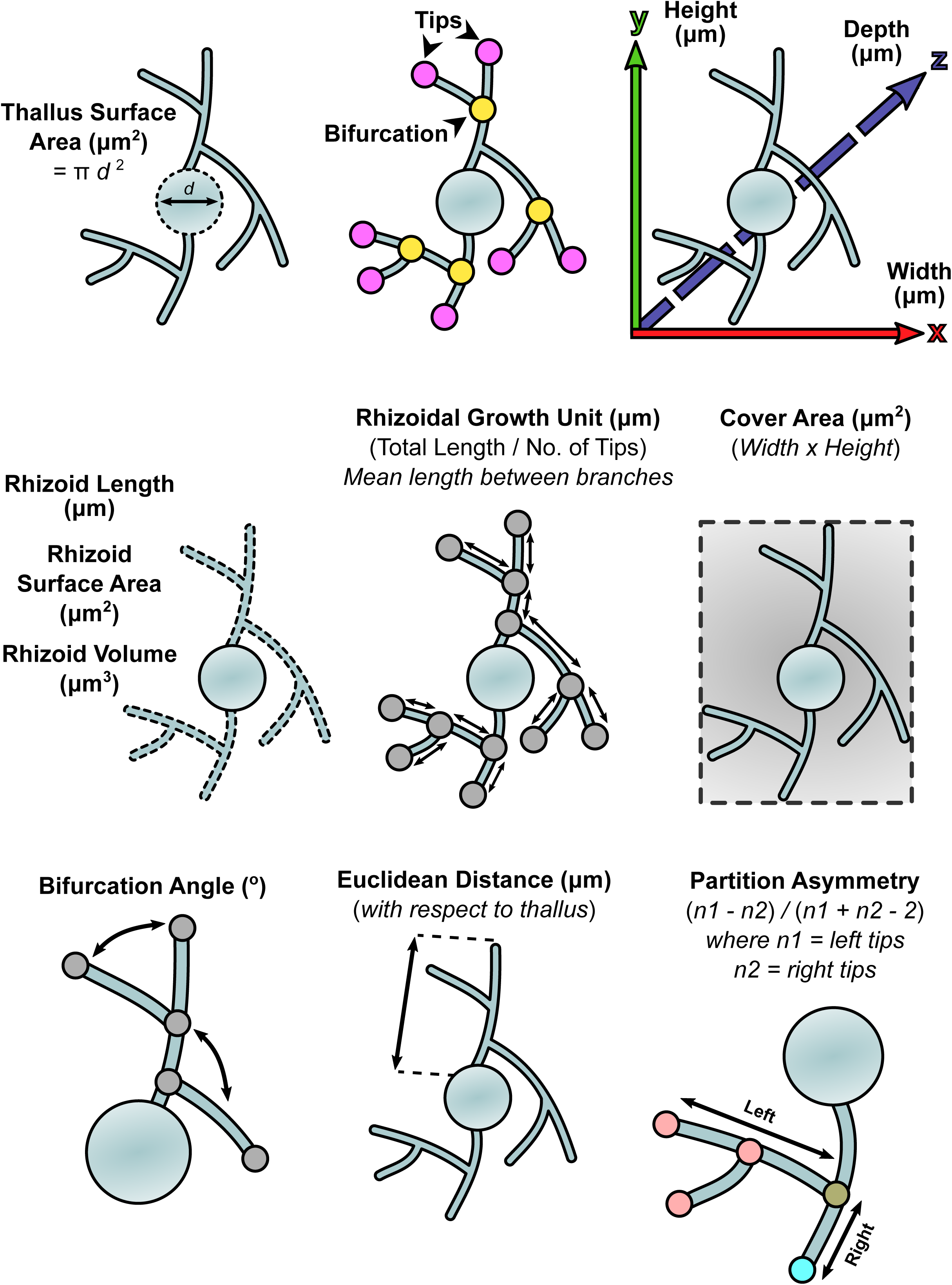
Chytrid rhizoids were quantified using morphometric parameters adapted from neurobiology. Diagrammatic glossary of neuronal morphometric parameters used to describe 3D reconstructed chytrid rhizoids from growth experiments. Chytrids are represented by an aerial 2D diagram, as if from a z-stack maximum intensity projection.

**Supplementary Figure 6.**
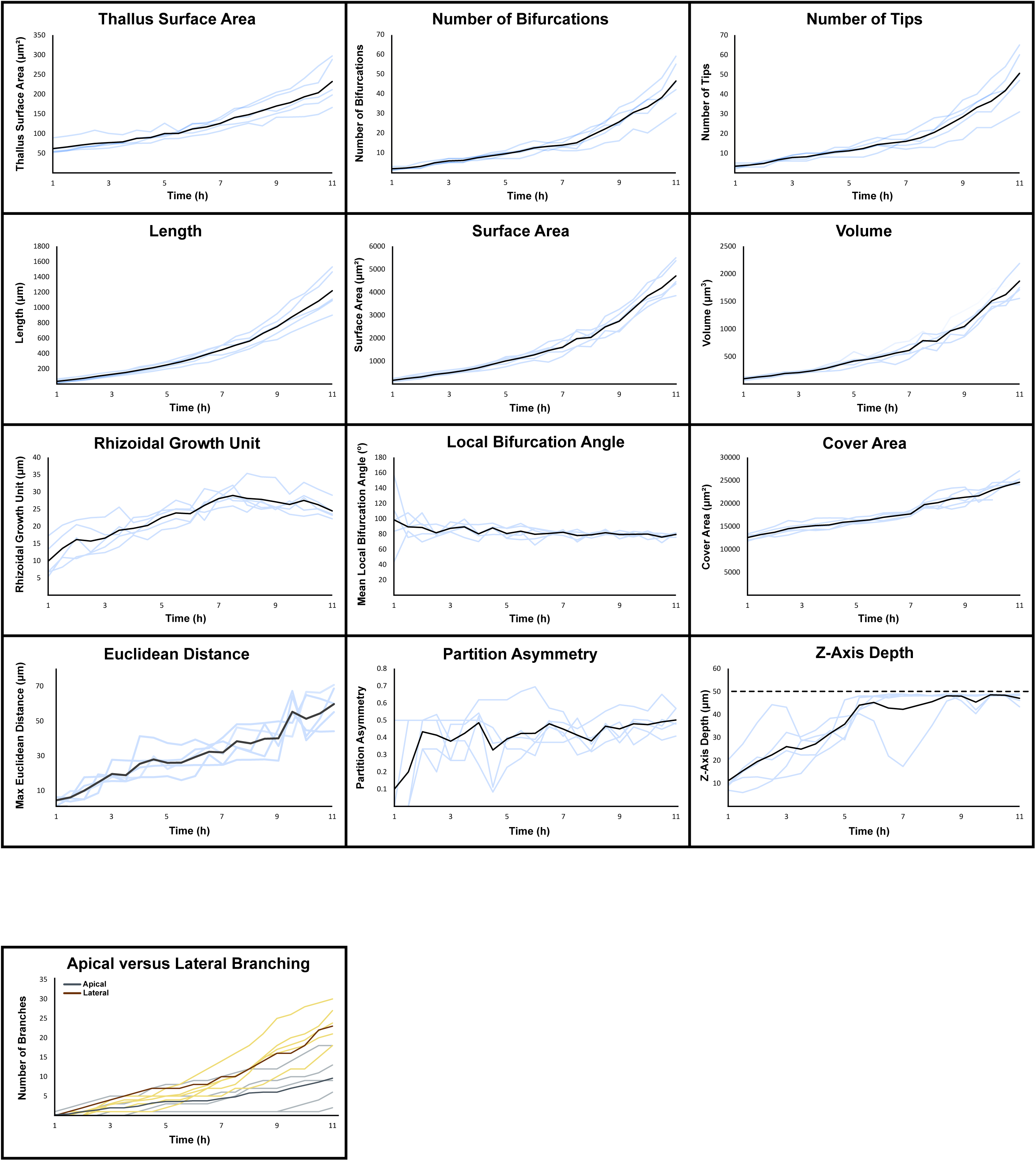
Development trajectories of major morphometric traits in *R. globosum* rhizoids. Growth patterns of morphometric features for developing *R. globosum* rhizoids taken from 4D microscopy experiments. Plateau in the z-axis depth occurs due growth outside of the designated experimental imaging field. Scale bar = 20 µm.

**Supplementary Figure 7.**
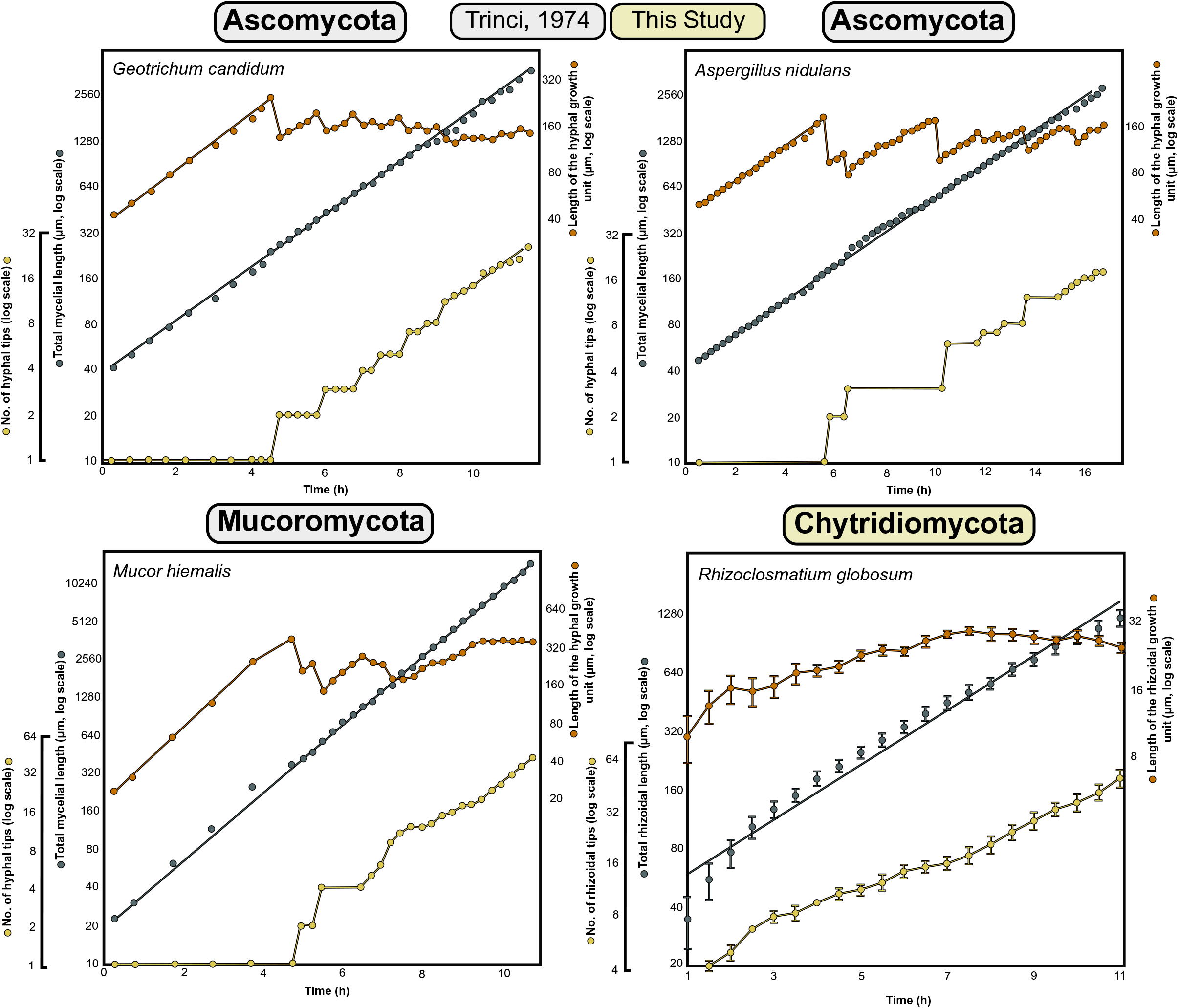
Development of chytrid rhizoids fundamentally resembles mycelial development in hyphal fungi. Comparison of the growth trajectories of the growth unit, total length and number of tips of the rhizoids or hyphae in fungi from the Ascomycota, Basidiomycota, Mucoromycota and Chytridiomycota. Data for Ascomycota, Basidiomycota and Mucoromycota fungi are not from this study and are reproduced as new figures directly from (Trinci, 1974).

**Supplementary Figure 8.**
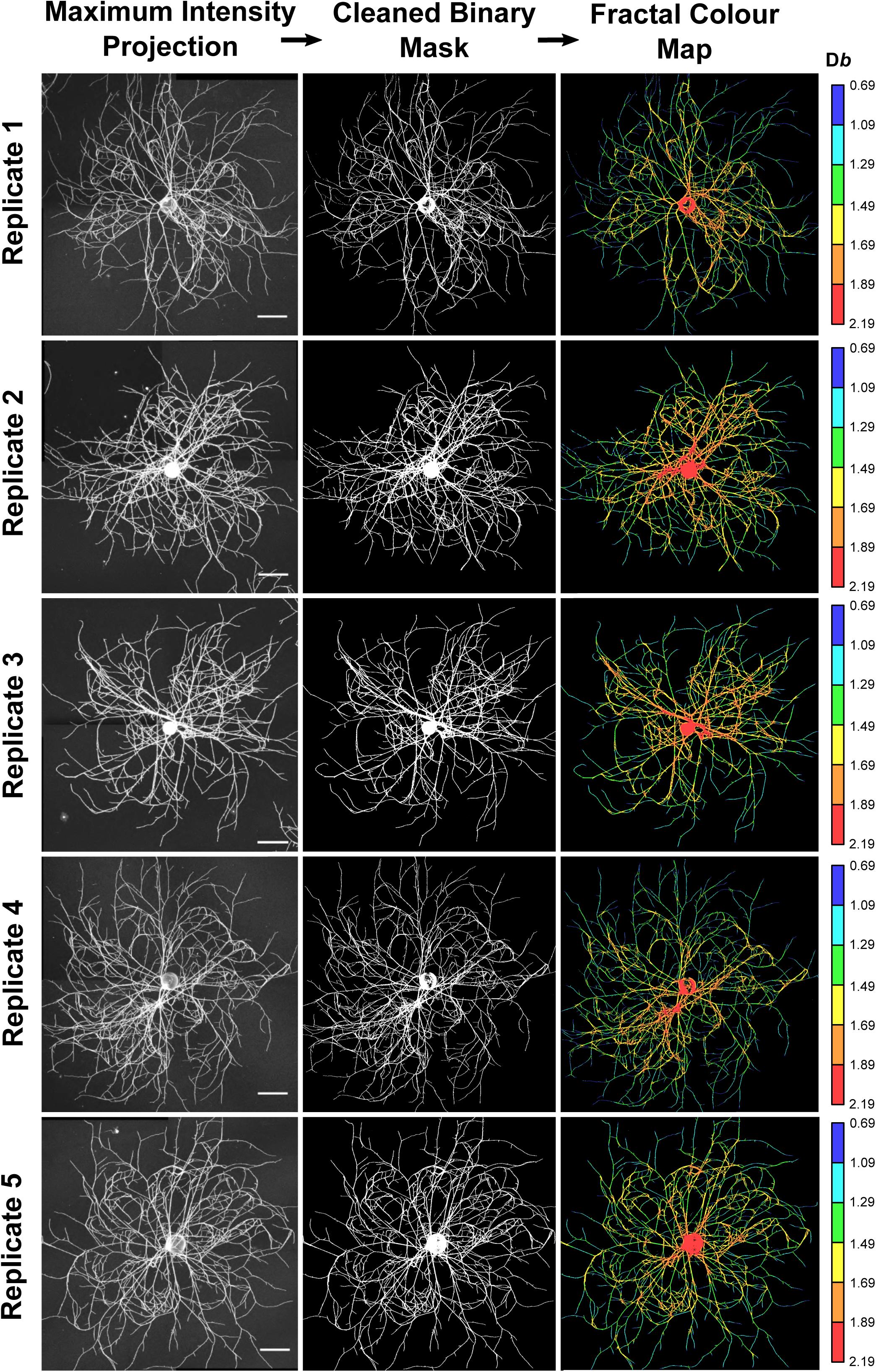
Fractal organisation of the chytrid rhizoid resembles that of mycelial colonies. Processing and fractal analysis workflow for 24 h *R. globosum* cells. Chytrid rhizoid systems become decreasingly fractal towards the growing edge. Final column images are pseudo-coloured by fractal dimension.

**Supplementary Table 1.** Morphometric features of developing *R. globosum* rhizoids associated with Figure 1 E-G.

| **4D Development** |  |  |  |  |  |  |
| --- | --- | --- | --- | --- | --- | --- |
| **Morphometric Feature** | **(1 h)**<br>**(Mean)** | **±**<br>**Standard**<br>**Deviation** | **(1.5 h)**<br>**(Mean)** | **±**<br>**Standard**<br>**Deviation** | **(2 h)**<br>**(Mean)** | **±**<br>**Standard**<br>**Deviation** |
| Thallus Diameter ($\mu\text{m}^2$) | 61.73 | 15.55 | 65.73 | 15.90 | 70.65 | 15.91 |
| Number of Bifurcations | 1.80 | 0.84 | 2.20 | 0.45 | 3.00 | 1.22 |
| Number of Tips | 3.40 | 1.14 | 4.00 | 0.71 | 4.80 | 1.10 |
| Width ($\mu\text{m}$) | 115.28 | 2.31 | 118.40 | 0.76 | 120.92 | 1.52 |
| Height ($\mu\text{m}$) | 108.89 | 4.02 | 111.68 | 5.43 | 113.31 | 7.81 |
| Depth ($\mu\text{m}$) | 11.13 | 5.23 | 15.48 | 7.68 | 19.48 | 10.83 |
| Total Length ($\mu\text{m}$) | 34.96 | 23.27 | 55.49 | 26.18 | 76.24 | 27.57 |
| Surface Area ($\mu\text{m}^2$) | 160.28 | 67.69 | 242.43 | 86.65 | 309.83 | 95.17 |
| Volume ($\mu\text{m}^3$) | 95.35 | 24.87 | 126.35 | 24.33 | 150.28 | 27.20 |
| Rhizoidal Growth Unit ($\mu\text{m}$) | 9.87 | 5.15 | 13.57 | 5.02 | 16.16 | 5.17 |
| Cover Area ($\mu\text{m}^2$) | 12558.93 | 680.10 | 13222.81 | 663.30 | 13701.19 | 945.32 |
| Mean Bifurcation Angle ($^\circ$) | 98.56 | 41.45 | 89.28 | 11.85 | 88.46 | 14.35 |
| Partition Asymmetry | 0.10 | 0.22 | 0.20 | 0.27 | 0.43 | 0.09 |
| Max Euclidean Distance ($\mu\text{m}$) | 4.59 | 2.06 | 6.24 | 2.21 | 10.10 | 5.36 |
| Max Path Distance ($\mu\text{m}$) | 4.92 | 2.43 | 7.21 | 2.93 | 11.50 | 5.88 |
| Number of Apical Branches | 0.20 | 0.45 | 0.40 | 0.89 | 1.00 | 1.22 |
| Number of Lateral Branches | 0.00 | 0.00 | 0.40 | 0.55 | 0.80 | 0.84 |
|  | **(2.5 h)**<br>**(Mean)** | **±**<br>**Standard**<br>**Deviation** | **(3 h)**<br>**(Mean)** | **±**<br>**Standard**<br>**Deviation** | **(3.5 h)**<br>**(Mean)** | **±**<br>**Standard**<br>**Deviation** |
| Thallus Diameter ($\mu\text{m}^2$) | 74.64 | 19.18 | 80.28 | 13.97 | 78.74 | 10.72 |
| Number of Bifurcations | 4.80 | 0.84 | 6.00 | 1.10 | 6.00 | 1.00 |
| Number of Tips | 6.60 | 0.55 | 8.17 | 1.30 | 8.20 | 1.79 |
| Width ($\mu\text{m}$) | 124.56 | 1.06 | 126.69 | 2.62 | 127.67 | 2.68 |
| Height ($\mu\text{m}$) | 116.33 | 9.44 | 117.53 | 8.31 | 118.83 | 10.80 |
| Depth ($\mu\text{m}$) | 22.43 | 13.50 | 24.05 | 12.05 | 24.90 | 6.66 |
| Total Length ($\mu\text{m}$) | 103.01 | 29.69 | 129.64 | 28.92 | 150.42 | 26.09 |
| Surface Area ($\mu\text{m}^2$) | 419.67 | 78.87 | 498.16 | 80.13 | 579.40 | 69.78 |
| Volume ($\mu\text{m}^3$) | 191.82 | 25.09 | 216.88 | 19.13 | 239.34 | 23.15 |
| Rhizoidal Growth Unit ($\mu\text{m}$) | 15.72 | 4.89 | 16.14 | 3.97 | 18.85 | 4.29 |
| Cover Area ($\mu\text{m}^2$) | 14489.81 | 1178.25 | 14876.19 | 798.19 | 15148.16 | 1056.39 |
| Mean Bifurcation Angle ($^\circ$) | 81.51 | 6.51 | 85.40 | 5.20 | 89.34 | 8.82 |
| Partition Asymmetry | 0.41 | 0.14 | 0.39 | 0.11 | 0.42 | 0.10 |
| Max Euclidean Distance ($\mu\text{m}$) | 14.85 | 4.23 | 19.28 | 4.71 | 18.90 | 4.97 |
| Max Path Distance ($\mu\text{m}$) | 18.14 | 5.13 | 23.20 | 5.13 | 23.75 | 5.18 |
| Number of Apical Branches | 1.40 | 1.67 | 2.50 | 2.00 | 2.00 | 2.00 |
| Number of Lateral Branches | 1.80 | 0.84 | 3.00 | 1.30 | 3.40 | 1.82 |
|  | **(4 h)**<br>**(Mean)** | **±**<br>**Standard**<br>**Deviation** | **(4.5 h)**<br>**(Mean)** | **±**<br>**Standard**<br>**Deviation** | **(5 h)**<br>**(Mean)** | **±**<br>**Standard**<br>**Deviation** |
| Thallus Diameter ($\mu\text{m}^2$) | 87.55 | 13.18 | 89.84 | 10.18 | 99.90 | 15.33 |
| Number of Bifurcations | 7.40 | 0.89 | 8.40 | 1.34 | 9.40 | 1.52 |
| Number of Tips | 9.40 | 0.89 | 10.60 | 1.82 | 11.20 | 1.92 |
| Width ($\mu\text{m}$) | 128.49 | 4.98 | 130.81 | 6.80 | 133.09 | 8.08 |
| Height ($\mu\text{m}$) | 119.48 | 13.29 | 121.85 | 11.97 | 121.57 | 10.10 |
| Depth ($\mu\text{m}$) | 27.16 | 5.01 | 31.81 | 4.67 | 35.82 | 7.13 |
| Total Length ($\mu\text{m}$) | 183.21 | 34.22 | 212.60 | 35.17 | 250.30 | 39.92 |
| Surface Area ($\mu\text{m}^2$) | 697.97 | 112.24 | 856.05 | 134.33 | 1004.34 | 184.45 |
| Volume ($\mu\text{m}^3$) | 286.28 | 51.44 | 351.77 | 63.07 | 416.94 | 108.92 |
| Rhizoidal Growth Unit ($\mu\text{m}$) | 19.40 | 2.28 | 20.23 | 2.71 | 22.54 | 2.55 |
| Cover Area ($\mu\text{m}^2$) | 15298.59 | 1096.41 | 15874.79 | 707.32 | 16119.89 | 603.53 |
| Mean Bifurcation Angle ($^\circ$) | 80.11 | 7.99 | 87.90 | 5.39 | 80.59 | 6.54 |
| Partition Asymmetry | 0.49 | 0.14 | 0.33 | 0.23 | 0.39 | 0.14 |
| Max Euclidean Distance ($\mu\text{m}$) | 25.42 | 9.73 | 27.94 | 8.30 | 26.01 | 7.13 |
| Max Path Distance ($\mu\text{m}$) | 30.92 | 11.74 | 35.03 | 10.52 | 34.40 | 11.86 |
| Number of Apical Branches | 2.40 | 1.95 | 3.20 | 2.68 | 3.60 | 2.97 |
| Number of Lateral Branches | 3.80 | 1.92 | 4.20 | 2.28 | 5.00 | 2.12 |
|  | **(5.5 h)**<br>**(Mean)** | **±**<br>**Standard**<br>**Deviation** | **(6 h)**<br>**(Mean)** | **±**<br>**Standard**<br>**Deviation** | **(6.5 h)**<br>**(Mean)** | **±**<br>**Standard**<br>**Deviation** |
| Thallus Diameter ($\mu\text{m}^2$) | 100.34 | 7.52 | 111.63 | 13.83 | 116.66 | 12.51 |
| Number of Bifurcations | 10.60 | 2.51 | 12.40 | 2.51 | 13.20 | 1.79 |
| Number of Tips | 12.40 | 2.97 | 14.40 | 2.97 | 15.20 | 2.68 |
| Width ($\mu\text{m}$) | 134.75 | 8.19 | 137.86 | 8.19 | 141.14 | 9.67 |
| Height ($\mu\text{m}$) | 122.19 | 8.54 | 123.33 | 8.78 | 123.15 | 8.15 |
| Depth ($\mu\text{m}$) | 44.05 | 3.44 | 45.24 | 4.66 | 42.83 | 11.70 |
| Total Length ($\mu\text{m}$) | 290.01 | 48.65 | 336.86 | 55.51 | 394.34 | 70.39 |
| Surface Area ($\mu\text{m}^2$) | 1131.95 | 138.18 | 1268.24 | 186.91 | 1460.13 | 325.84 |
| Volume ($\mu\text{m}^3$) | 448.43 | 45.57 | 497.52 | 79.06 | 560.30 | 140.57 |
| Rhizoidal Growth Unit ($\mu\text{m}$) | 23.87 | 3.07 | 23.65 | 2.33 | 26.07 | 3.45 |
| Cover Area ($\mu\text{m}^2$) | 16414.85 | 525.82 | 16960.99 | 817.17 | 17329.43 | 639.39 |
| Mean Bifurcation Angle ($^\circ$) | 83.58 | 8.62 | 79.63 | 9.66 | 80.75 | 2.71 |
| Partition Asymmetry | 0.42 | 0.15 | 0.42 | 0.16 | 0.48 | 0.07 |
| Max Euclidean Distance ($\mu\text{m}$) | 26.12 | 7.24 | 29.34 | 8.08 | 32.36 | 4.80 |
| Max Path Distance ($\mu\text{m}$) | 35.29 | 10.43 | 39.00 | 12.77 | 44.57 | 8.77 |
| Number of Apical Branches | 3.60 | 2.97 | 3.80 | 3.35 | 3.80 | 3.35 |
| Number of Lateral Branches | 5.40 | 2.07 | 6.80 | 2.17 | 7.80 | 2.59 |
|  | **(7 h)**<br>**(Mean)** | **±**<br>**Standard**<br>**Deviation** | **(7.5 h)**<br>**(Mean)** | **±**<br>**Standard**<br>**Deviation** | **(8 h)**<br>**(Mean)** | **±**<br>**Standard**<br>**Deviation** |
| Thallus Diameter ($\mu\text{m}^2$) | 125.80 | 13.55 | 140.79 | 20.30 | 153.26 | 21.19 |
| Number of Bifurcations | 13.80 | 2.28 | 15.00 | 3.81 | 19.83 | 4.28 |
| Number of Tips | 16.00 | 3.08 | 17.80 | 4.55 | 22.00 | 5.55 |
| Width ($\mu\text{m}$) | 144.14 | 10.96 | 149.97 | 9.68 | 150.13 | 11.91 |
| Height ($\mu\text{m}$) | 123.26 | 6.90 | 131.91 | 7.67 | 136.32 | 9.30 |
| Depth ($\mu\text{m}$) | 42.26 | 13.92 | 43.90 | 9.92 | 46.12 | 5.77 |
| Total Length ($\mu\text{m}$) | 447.68 | 83.46 | 507.96 | 93.67 | 590.88 | 85.01 |
| Surface Area ($\mu\text{m}^2$) | 1602.52 | 321.61 | 1977.68 | 343.82 | 2181.31 | 275.48 |
| Volume ($\mu\text{m}^3$) | 609.97 | 132.73 | 786.87 | 146.15 | 847.84 | 112.19 |
| Rhizoidal Growth Unit ($\mu\text{m}$) | 28.05 | 1.97 | 28.96 | 2.52 | 27.66 | 4.29 |
| Cover Area ($\mu\text{m}^2$) | 17710.48 | 522.73 | 19733.66 | 687.82 | 20439.86 | 1649.74 |
| Mean Bifurcation Angle ($^\circ$) | 82.27 | 2.76 | 77.85 | 6.87 | 78.96 | 2.85 |
| Partition Asymmetry | 0.45 | 0.07 | 0.41 | 0.08 | 0.40 | 0.07 |
| Max Euclidean Distance ($\mu\text{m}$) | 31.95 | 7.37 | 38.37 | 8.62 | 38.31 | 8.82 |
| Max Path Distance ($\mu\text{m}$) | 44.88 | 11.89 | 53.08 | 14.99 | 55.74 | 13.07 |
| Number of Apical Branches | 4.40 | 3.78 | 4.80 | 4.15 | 6.83 | 4.76 |
| Number of Lateral Branches | 9.00 | 3.39 | 10.20 | 3.49 | 12.50 | 3.94 |

|  | **(8.5 h)**<br>**(Mean)** | **±**<br>**Standard**<br>**Deviation** | **(9 h)**<br>**(Mean)** | **±**<br>**Standard**<br>**Deviation** | **(9.5 h)**<br>**(Mean)** | **±**<br>**Standard**<br>**Deviation** |
| --- | --- | --- | --- | --- | --- | --- |
| Thallus Diameter ($\mu\text{m}^2$) | 158.24 | 28.84 | 169.98 | 28.77 | 178.45 | 30.65 |
| Number of Bifurcations | 22.00 | 4.53 | 25.60 | 6.50 | 30.40 | 5.18 |
| Number of Tips | 24.40 | 5.73 | 28.40 | 7.50 | 33.20 | 6.53 |
| Width ($\mu\text{m}$) | 153.49 | 15.50 | 154.51 | 15.11 | 157.71 | 17.56 |
| Height ($\mu\text{m}$) | 136.95 | 9.97 | 138.78 | 14.09 | 137.92 | 9.11 |
| Depth ($\mu\text{m}$) | 48.11 | 1.32 | 48.03 | 1.18 | 45.40 | 3.89 |
| Total Length ($\mu\text{m}$) | 664.75 | 106.31 | 749.34 | 131.57 | 869.09 | 162.80 |
| Surface Area ($\mu\text{m}^2$) | 2484.50 | 341.81 | 2736.00 | 427.52 | 3290.47 | 372.99 |
| Volume ($\mu\text{m}^3$) | 963.75 | 163.46 | 1043.59 | 195.90 | 1269.05 | 150.26 |
| Rhizoidal Growth Unit ($\mu\text{m}$) | 27.84 | 3.78 | 27.12 | 3.95 | 26.35 | 2.25 |
| Cover Area ($\mu\text{m}^2$) | 20955.39 | 1838.27 | 21331.96 | 1840.64 | 21626.97 | 1102.86 |
| Mean Bifurcation Angle ($^\circ$) | 81.69 | 3.23 | 79.24 | 4.05 | 79.52 | 2.78 |
| Partition Asymmetry | 0.47 | 0.06 | 0.45 | 0.09 | 0.48 | 0.08 |
| Max Euclidean Distance ($\mu\text{m}$) | 39.80 | 10.19 | 40.08 | 8.86 | 55.31 | 10.81 |
| Max Path Distance ($\mu\text{m}$) | 55.92 | 12.69 | 57.55 | 16.95 | 74.23 | 12.13 |
| Number of Apical Branches | 6.00 | 4.90 | 6.00 | 4.90 | 7.00 | 5.48 |
| Number of Lateral Branches | 14.60 | 4.62 | 17.00 | 5.39 | 18.20 | 5.22 |
|  | **(10 h)**<br>**(Mean)** | **±**<br>**Standard**<br>**Deviation** | **(10.5 h)**<br>**(Mean)** | **±**<br>**Standard**<br>**Deviation** | **(11 h)**<br>**(Mean)** | **±**<br>**Standard**<br>**Deviation** |
| Thallus Diameter ($\mu\text{m}^2$) | 193.39 | 38.51 | 203.54 | 47.69 | 232.18 | 57.64 |
| Number of Bifurcations | 33.20 | 8.53 | 38.00 | 8.46 | 46.40 | 11.41 |
| Number of Tips | 36.40 | 9.61 | 41.80 | 10.03 | 50.60 | 13.16 |
| Width ($\mu\text{m}$) | 166.25 | 21.27 | 168.86 | 22.61 | 170.74 | 25.31 |
| Height ($\mu\text{m}$) | 138.76 | 12.08 | 142.45 | 15.78 | 145.67 | 16.12 |
| Depth ($\mu\text{m}$) | 48.55 | 0.42 | 48.39 | 0.30 | 47.04 | 2.34 |
| Total Length ($\mu\text{m}$) | 977.64 | 186.20 | 1084.53 | 220.20 | 1219.15 | 268.22 |
| Surface Area ($\mu\text{m}^2$) | 3836.65 | 422.94 | 4187.25 | 549.42 | 4707.64 | 704.53 |
| Volume ($\mu\text{m}^3$) | 1510.22 | 145.44 | 1623.77 | 209.37 | 1868.34 | 280.25 |
| Rhizoidal Growth Unit ($\mu\text{m}$) | 27.51 | 3.84 | 26.30 | 2.78 | 24.48 | 2.67 |
| Cover Area ($\mu\text{m}^2$) | 22884.73 | 1500.84 | 23811.10 | 1536.94 | 24595.64 | 1879.28 |
| Mean Bifurcation Angle ($^\circ$) | 79.91 | 4.24 | 75.90 | 4.53 | 79.41 | 2.45 |
| Partition Asymmetry | 0.47 | 0.06 | 0.49 | 0.10 | 0.50 | 0.07 |
| Max Euclidean Distance ($\mu\text{m}$) | 51.27 | 13.40 | 54.37 | 9.71 | 59.69 | 10.78 |
| Max Path Distance ($\mu\text{m}$) | 72.72 | 13.07 | 78.64 | 11.26 | 86.12 | 12.58 |
| Number of Apical Branches | 7.80 | 5.97 | 8.60 | 6.58 | 9.60 | 6.19 |
| Number of Lateral Branches | 19.40 | 5.81 | 21.80 | 5.07 | 23.80 | 4.76 |

**Supplementary Table 2.**
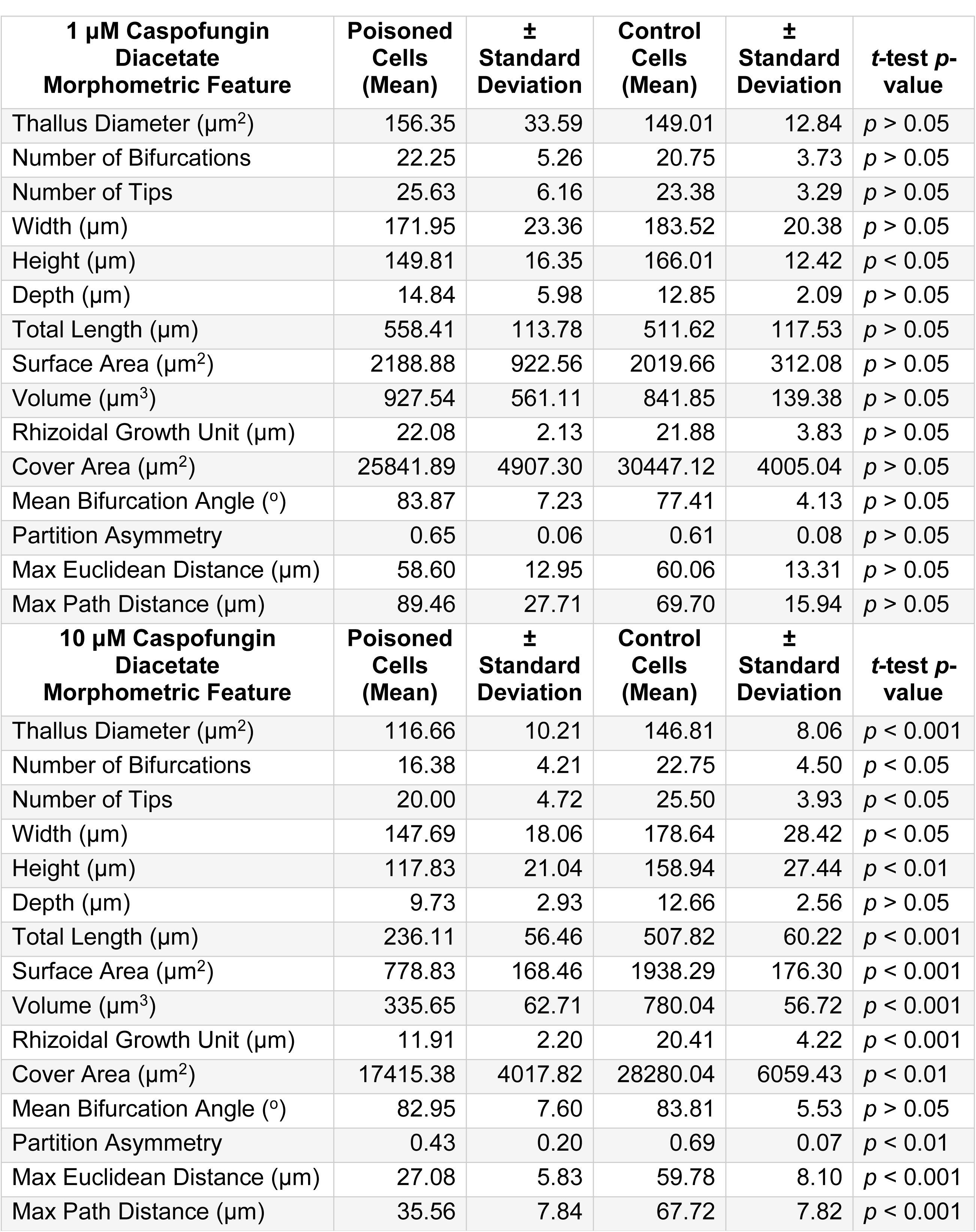
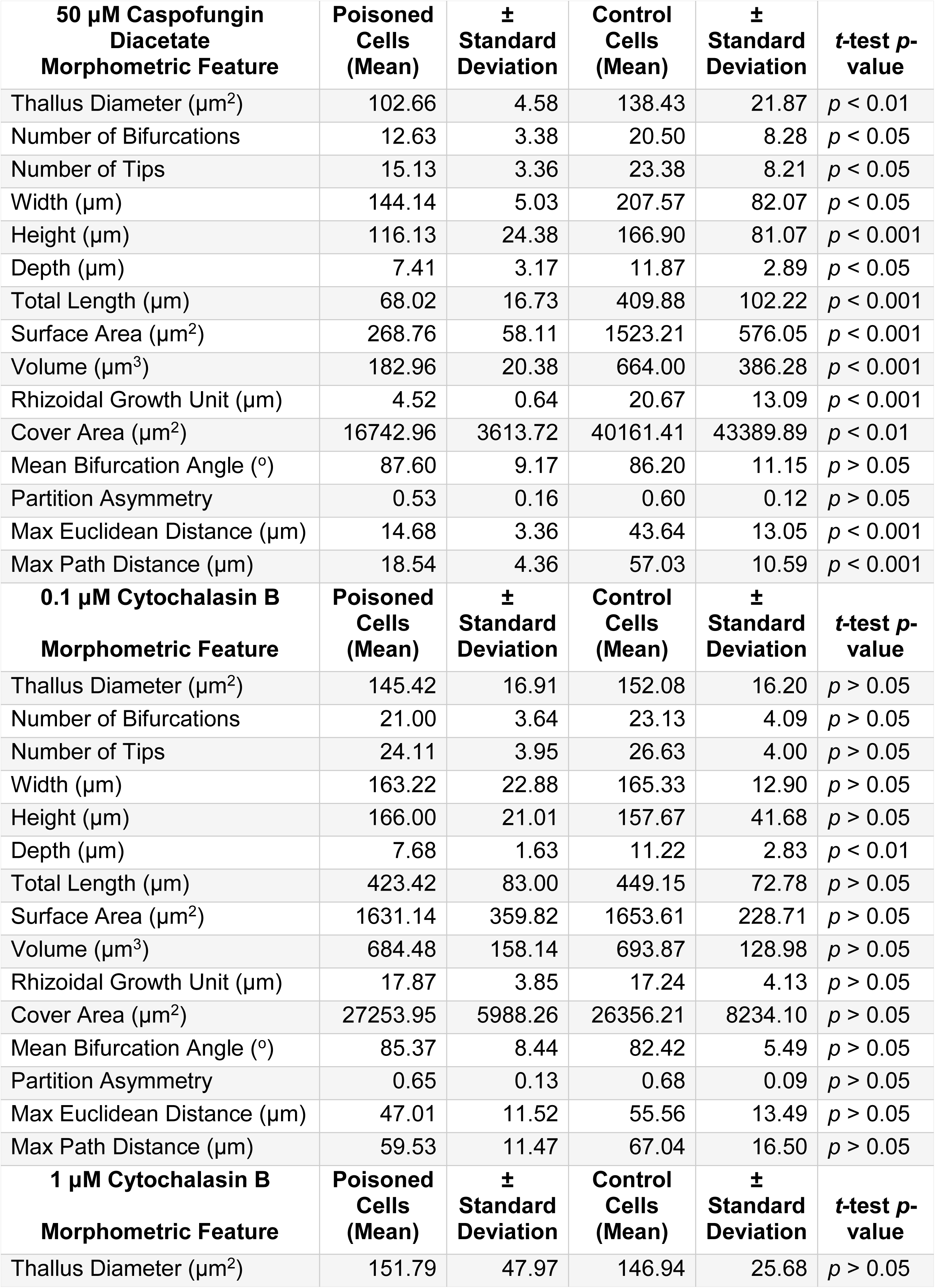
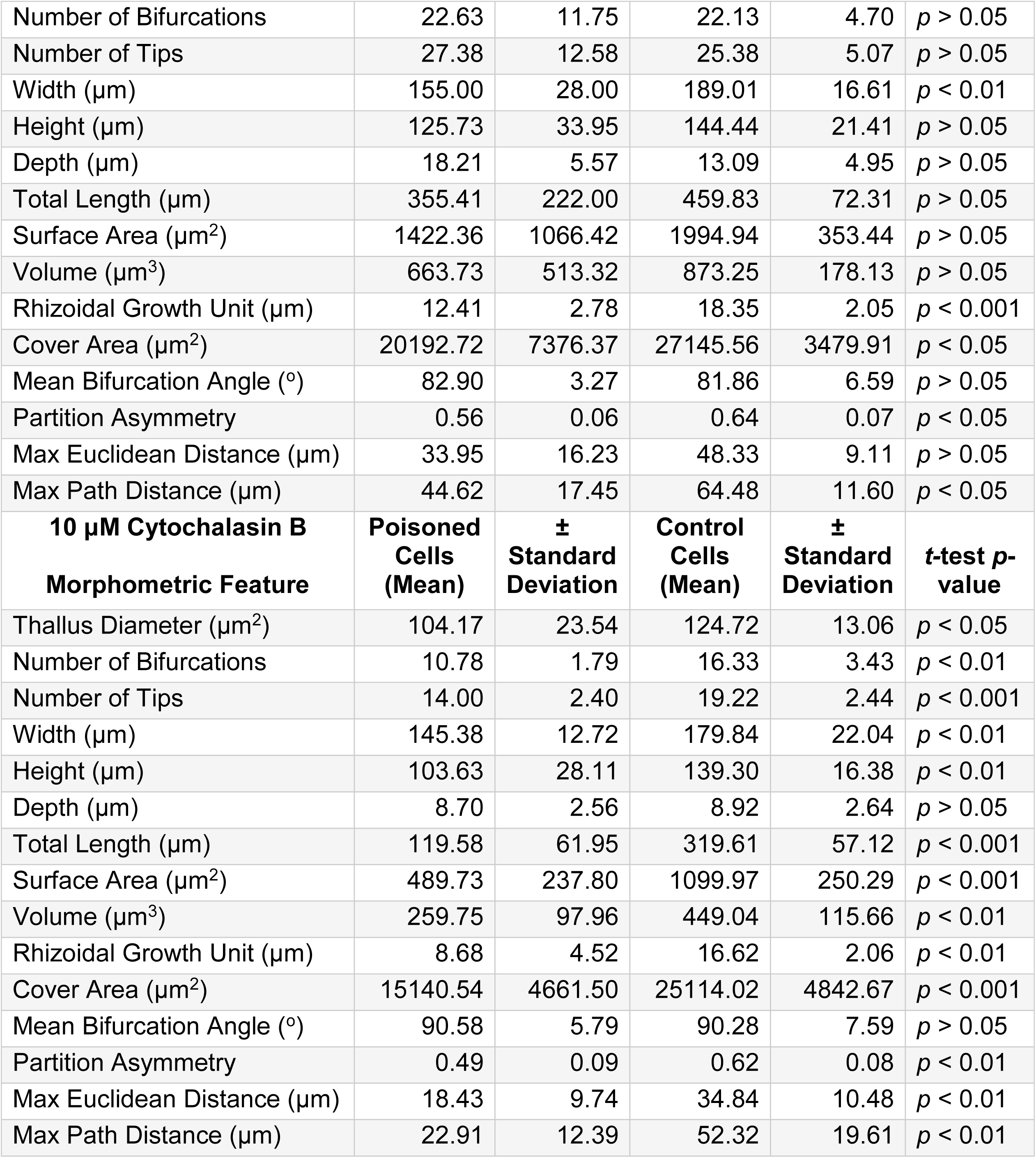
Morphometric features and statistical comparisons of chemically inhibited *R. globosum* rhizoids, associated with Figure 2 B-C.

| **1 μM Caspofungin Diacetate Morphometric Feature** | **Poisoned Cells (Mean)** | **± Standard Deviation** | **Control Cells (Mean)** | **± Standard Deviation** | **t-test p-value** |
| --- | --- | --- | --- | --- | --- |
| Thallus Diameter ($\mu\text{m}^2$) | 156.35 | 33.59 | 149.01 | 12.84 | $p > 0.05$ |
| Number of Bifurcations | 22.25 | 5.26 | 20.75 | 3.73 | $p > 0.05$ |
| Number of Tips | 25.63 | 6.16 | 23.38 | 3.29 | $p > 0.05$ |
| Width ($\mu\text{m}$) | 171.95 | 23.36 | 183.52 | 20.38 | $p > 0.05$ |
| Height ($\mu\text{m}$) | 149.81 | 16.35 | 166.01 | 12.42 | $p < 0.05$ |
| Depth ($\mu\text{m}$) | 14.84 | 5.98 | 12.85 | 2.09 | $p > 0.05$ |
| Total Length ($\mu\text{m}$) | 558.41 | 113.78 | 511.62 | 117.53 | $p > 0.05$ |
| Surface Area ($\mu\text{m}^2$) | 2188.88 | 922.56 | 2019.66 | 312.08 | $p > 0.05$ |
| Volume ($\mu\text{m}^3$) | 927.54 | 561.11 | 841.85 | 139.38 | $p > 0.05$ |
| Rhizoidal Growth Unit ($\mu\text{m}$) | 22.08 | 2.13 | 21.88 | 3.83 | $p > 0.05$ |
| Cover Area ($\mu\text{m}^2$) | 25841.89 | 4907.30 | 30447.12 | 4005.04 | $p > 0.05$ |
| Mean Bifurcation Angle ($^\circ$) | 83.87 | 7.23 | 77.41 | 4.13 | $p > 0.05$ |
| Partition Asymmetry | 0.65 | 0.06 | 0.61 | 0.08 | $p > 0.05$ |
| Max Euclidean Distance ($\mu\text{m}$) | 58.60 | 12.95 | 60.06 | 13.31 | $p > 0.05$ |
| Max Path Distance ($\mu\text{m}$) | 89.46 | 27.71 | 69.70 | 15.94 | $p > 0.05$ |
| **10 μM Caspofungin Diacetate Morphometric Feature** | **Poisoned Cells (Mean)** | **± Standard Deviation** | **Control Cells (Mean)** | **± Standard Deviation** | **t-test p-value** |
| Thallus Diameter ($\mu\text{m}^2$) | 116.66 | 10.21 | 146.81 | 8.06 | $p < 0.001$ |
| Number of Bifurcations | 16.38 | 4.21 | 22.75 | 4.50 | $p < 0.05$ |
| Number of Tips | 20.00 | 4.72 | 25.50 | 3.93 | $p < 0.05$ |
| Width ($\mu\text{m}$) | 147.69 | 18.06 | 178.64 | 28.42 | $p < 0.05$ |
| Height ($\mu\text{m}$) | 117.83 | 21.04 | 158.94 | 27.44 | $p < 0.01$ |
| Depth ($\mu\text{m}$) | 9.73 | 2.93 | 12.66 | 2.56 | $p > 0.05$ |
| Total Length ($\mu\text{m}$) | 236.11 | 56.46 | 507.82 | 60.22 | $p < 0.001$ |
| Surface Area ($\mu\text{m}^2$) | 778.83 | 168.46 | 1938.29 | 176.30 | $p < 0.001$ |
| Volume ($\mu\text{m}^3$) | 335.65 | 62.71 | 780.04 | 56.72 | $p < 0.001$ |
| Rhizoidal Growth Unit ($\mu\text{m}$) | 11.91 | 2.20 | 20.41 | 4.22 | $p < 0.001$ |
| Cover Area ($\mu\text{m}^2$) | 17415.38 | 4017.82 | 28280.04 | 6059.43 | $p < 0.01$ |
| Mean Bifurcation Angle ($^\circ$) | 82.95 | 7.60 | 83.81 | 5.53 | $p > 0.05$ |
| Partition Asymmetry | 0.43 | 0.20 | 0.69 | 0.07 | $p < 0.01$ |
| Max Euclidean Distance ($\mu\text{m}$) | 27.08 | 5.83 | 59.78 | 8.10 | $p < 0.001$ |
| Max Path Distance ($\mu\text{m}$) | 35.56 | 7.84 | 67.72 | 7.82 | $p < 0.001$ |

| **50 μM Caspofungin Diacetate Morphometric Feature** | **Poisoned Cells (Mean)** | **± Standard Deviation** | **Control Cells (Mean)** | **± Standard Deviation** | ***t*-test *p*-value** |
| --- | --- | --- | --- | --- | --- |
| Thallus Diameter ($\mu\text{m}^2$) | 102.66 | 4.58 | 138.43 | 21.87 | $p < 0.01$ |
| Number of Bifurcations | 12.63 | 3.38 | 20.50 | 8.28 | $p < 0.05$ |
| Number of Tips | 15.13 | 3.36 | 23.38 | 8.21 | $p < 0.05$ |
| Width ($\mu\text{m}$) | 144.14 | 5.03 | 207.57 | 82.07 | $p < 0.05$ |
| Height ($\mu\text{m}$) | 116.13 | 24.38 | 166.90 | 81.07 | $p < 0.001$ |
| Depth ($\mu\text{m}$) | 7.41 | 3.17 | 11.87 | 2.89 | $p < 0.05$ |
| Total Length ($\mu\text{m}$) | 68.02 | 16.73 | 409.88 | 102.22 | $p < 0.001$ |
| Surface Area ($\mu\text{m}^2$) | 268.76 | 58.11 | 1523.21 | 576.05 | $p < 0.001$ |
| Volume ($\mu\text{m}^3$) | 182.96 | 20.38 | 664.00 | 386.28 | $p < 0.001$ |
| Rhizoidal Growth Unit ($\mu\text{m}$) | 4.52 | 0.64 | 20.67 | 13.09 | $p < 0.001$ |
| Cover Area ($\mu\text{m}^2$) | 16742.96 | 3613.72 | 40161.41 | 43389.89 | $p < 0.01$ |
| Mean Bifurcation Angle ($^\circ$) | 87.60 | 9.17 | 86.20 | 11.15 | $p > 0.05$ |
| Partition Asymmetry | 0.53 | 0.16 | 0.60 | 0.12 | $p > 0.05$ |
| Max Euclidean Distance ($\mu\text{m}$) | 14.68 | 3.36 | 43.64 | 13.05 | $p < 0.001$ |
| Max Path Distance ($\mu\text{m}$) | 18.54 | 4.36 | 57.03 | 10.59 | $p < 0.001$ |
| **0.1 μM Cytochalasin B Morphometric Feature** | **Poisoned Cells (Mean)** | **± Standard Deviation** | **Control Cells (Mean)** | **± Standard Deviation** | ***t*-test *p*-value** |
| Thallus Diameter ($\mu\text{m}^2$) | 145.42 | 16.91 | 152.08 | 16.20 | $p > 0.05$ |
| Number of Bifurcations | 21.00 | 3.64 | 23.13 | 4.09 | $p > 0.05$ |
| Number of Tips | 24.11 | 3.95 | 26.63 | 4.00 | $p > 0.05$ |
| Width ($\mu\text{m}$) | 163.22 | 22.88 | 165.33 | 12.90 | $p > 0.05$ |
| Height ($\mu\text{m}$) | 166.00 | 21.01 | 157.67 | 41.68 | $p > 0.05$ |
| Depth ($\mu\text{m}$) | 7.68 | 1.63 | 11.22 | 2.83 | $p < 0.01$ |
| Total Length ($\mu\text{m}$) | 423.42 | 83.00 | 449.15 | 72.78 | $p > 0.05$ |
| Surface Area ($\mu\text{m}^2$) | 1631.14 | 359.82 | 1653.61 | 228.71 | $p > 0.05$ |
| Volume ($\mu\text{m}^3$) | 684.48 | 158.14 | 693.87 | 128.98 | $p > 0.05$ |
| Rhizoidal Growth Unit ($\mu\text{m}$) | 17.87 | 3.85 | 17.24 | 4.13 | $p > 0.05$ |
| Cover Area ($\mu\text{m}^2$) | 27253.95 | 5988.26 | 26356.21 | 8234.10 | $p > 0.05$ |
| Mean Bifurcation Angle ($^\circ$) | 85.37 | 8.44 | 82.42 | 5.49 | $p > 0.05$ |
| Partition Asymmetry | 0.65 | 0.13 | 0.68 | 0.09 | $p > 0.05$ |
| Max Euclidean Distance ($\mu\text{m}$) | 47.01 | 11.52 | 55.56 | 13.49 | $p > 0.05$ |
| Max Path Distance ($\mu\text{m}$) | 59.53 | 11.47 | 67.04 | 16.50 | $p > 0.05$ |
| **1 μM Cytochalasin B Morphometric Feature** | **Poisoned Cells (Mean)** | **± Standard Deviation** | **Control Cells (Mean)** | **± Standard Deviation** | ***t*-test *p*-value** |
| Thallus Diameter ($\mu\text{m}^2$) | 151.79 | 47.97 | 146.94 | 25.68 | $p > 0.05$ |
| Number of Bifurcations | 22.63 | 11.75 | 22.13 | 4.70 | $p > 0.05$ |
| Number of Tips | 27.38 | 12.58 | 25.38 | 5.07 | $p > 0.05$ |
| Width ($\mu\text{m}$) | 155.00 | 28.00 | 189.01 | 16.61 | $p < 0.01$ |
| Height ($\mu\text{m}$) | 125.73 | 33.95 | 144.44 | 21.41 | $p > 0.05$ |
| Depth ($\mu\text{m}$) | 18.21 | 5.57 | 13.09 | 4.95 | $p > 0.05$ |
| Total Length ($\mu\text{m}$) | 355.41 | 222.00 | 459.83 | 72.31 | $p > 0.05$ |
| Surface Area ($\mu\text{m}^2$) | 1422.36 | 1066.42 | 1994.94 | 353.44 | $p > 0.05$ |
| Volume ($\mu\text{m}^3$) | 663.73 | 513.32 | 873.25 | 178.13 | $p > 0.05$ |
| Rhizoidal Growth Unit ($\mu\text{m}$) | 12.41 | 2.78 | 18.35 | 2.05 | $p < 0.001$ |
| Cover Area ($\mu\text{m}^2$) | 20192.72 | 7376.37 | 27145.56 | 3479.91 | $p < 0.05$ |
| Mean Bifurcation Angle ($^\circ$) | 82.90 | 3.27 | 81.86 | 6.59 | $p > 0.05$ |
| Partition Asymmetry | 0.56 | 0.06 | 0.64 | 0.07 | $p < 0.05$ |
| Max Euclidean Distance ($\mu\text{m}$) | 33.95 | 16.23 | 48.33 | 9.11 | $p > 0.05$ |
| Max Path Distance ($\mu\text{m}$) | 44.62 | 17.45 | 64.48 | 11.60 | $p < 0.05$ |
| **10 $\mu\text{M}$ Cytochalasin B** | **Poisoned** | **±** | **Control** | **±** |  |
| **Morphometric Feature** | **Cells** | **Standard** | **Cells** | **Standard** | **t-test p-value** |
|  | **(Mean)** | **Deviation** | **(Mean)** | **Deviation** |  |
| Thallus Diameter ($\mu\text{m}^2$) | 104.17 | 23.54 | 124.72 | 13.06 | $p < 0.05$ |
| Number of Bifurcations | 10.78 | 1.79 | 16.33 | 3.43 | $p < 0.01$ |
| Number of Tips | 14.00 | 2.40 | 19.22 | 2.44 | $p < 0.001$ |
| Width ($\mu\text{m}$) | 145.38 | 12.72 | 179.84 | 22.04 | $p < 0.01$ |
| Height ($\mu\text{m}$) | 103.63 | 28.11 | 139.30 | 16.38 | $p < 0.01$ |
| Depth ($\mu\text{m}$) | 8.70 | 2.56 | 8.92 | 2.64 | $p > 0.05$ |
| Total Length ($\mu\text{m}$) | 119.58 | 61.95 | 319.61 | 57.12 | $p < 0.001$ |
| Surface Area ($\mu\text{m}^2$) | 489.73 | 237.80 | 1099.97 | 250.29 | $p < 0.001$ |
| Volume ($\mu\text{m}^3$) | 259.75 | 97.96 | 449.04 | 115.66 | $p < 0.01$ |
| Rhizoidal Growth Unit ($\mu\text{m}$) | 8.68 | 4.52 | 16.62 | 2.06 | $p < 0.01$ |
| Cover Area ($\mu\text{m}^2$) | 15140.54 | 4661.50 | 25114.02 | 4842.67 | $p < 0.001$ |
| Mean Bifurcation Angle ($^\circ$) | 90.58 | 5.79 | 90.28 | 7.59 | $p > 0.05$ |
| Partition Asymmetry | 0.49 | 0.09 | 0.62 | 0.08 | $p < 0.01$ |
| Max Euclidean Distance ($\mu\text{m}$) | 18.43 | 9.74 | 34.84 | 10.48 | $p < 0.01$ |
| Max Path Distance ($\mu\text{m}$) | 22.91 | 12.39 | 52.32 | 19.61 | $p < 0.01$ |

**Supplementary Table 3.** Morphometric features and statistical comparisons of *R. globosum* rhizoids growing in carbon replete or deplete media, associated with Figure 3 A-C.

| **(1 h)**<br>**Morphometric Feature** | **Carbon**<br>**Replete**<br>**(Mean)** | **±**<br>**Standard**<br>**Deviation** | **Carbon**<br>**Deplete**<br>**(Mean)** | **±**<br>**Standard**<br>**Deviation** | ***t*-test *p*-**<br>**value** |
| --- | --- | --- | --- | --- | --- |
| Thallus Diameter (µm<sup>2</sup>) | 65.33 | 10.96 | 51.85 | 11.85 | *p* < 0.01 |
| Number of Bifurcations | 4.33 | 1.58 | 3.63 | 2.70 | *p* > 0.05 |
| Number of Tips | 5.78 | 1.30 | 4.75 | 2.60 | *p* > 0.05 |
| Width (µm) | 47.59 | 7.42 | 42.38 | 22.60 | *p* > 0.05 |
| Height (µm) | 49.04 | 12.71 | 45.21 | 20.65 | *p* > 0.05 |
| Depth (µm) | 7.58 | 2.60 | 6.74 | 1.68 | *p* > 0.05 |
| Total Length (µm) | 75.54 | 16.57 | 74.47 | 55.59 | *p* > 0.05 |
| Surface Area (µm<sup>2</sup>) | 207.65 | 36.34 | 173.69 | 205.38 | *p* > 0.05 |
| Volume (µm<sup>3</sup>) | 98.70 | 22.65 | 70.36 | 70.72 | *p* < 0.001 |
| Rhizoidal Growth Unit (µm) | 13.19 | 1.53 | 15.82 | 2.75 | *p* > 0.05 |
| Cover Area (µm<sup>2</sup>) | 2400.60 | 934.82 | 1959.75 | 2892.47 | *p* > 0.05 |
| Mean Bifurcation Angle (°) | 93.94 | 16.81 | 92.24 | 16.11 | *p* > 0.05 |
| Partition Asymmetry | 0.50 | 0.13 | 0.54 | 0.17 | *p* > 0.05 |
| Max Euclidean Distance (µm) | 11.26 | 4.81 | 13.08 | 11.37 | *p* > 0.05 |
| Max Path Distance (µm) | 13.67 | 5.78 | 16.05 | 12.60 | *p* > 0.05 |
| **(4 h)**<br>**Morphometric Feature** | **Carbon**<br>**Replete**<br>**(Mean)** | **±**<br>**Standard**<br>**Deviation** | **Carbon**<br>**Deplete**<br>**(Mean)** | **±**<br>**Standard**<br>**Deviation** | ***t*-test *p*-**<br>**value** |
| Thallus Diameter (µm<sup>2</sup>) | 103.87 | 16.89 | 70.28 | 10.53 | *p* < 0.001 |
| Number of Bifurcations | 14.38 | 4.47 | 10.44 | 2.88 | *p* > 0.05 |
| Number of Tips | 16.25 | 4.33 | 12.00 | 2.74 | *p* < 0.05 |
| Width (µm) | 102.31 | 17.57 | 142.60 | 20.37 | *p* < 0.001 |
| Height (µm) | 103.86 | 22.07 | 152.28 | 40.59 | *p* < 0.01 |
| Depth (µm) | 9.35 | 1.63 | 7.23 | 1.68 | *p* < 0.05 |
| Total Length (µm) | 297.44 | 91.96 | 400.84 | 41.77 | *p* < 0.05 |
| Surface Area (µm<sup>2</sup>) | 1052.25 | 355.81 | 1353.25 | 399.14 | *p* > 0.05 |
| Volume (µm<sup>3</sup>) | 413.31 | 174.59 | 440.55 | 231.56 | *p* > 0.05 |
| Rhizoidal Growth Unit (µm) | 18.23 | 2.12 | 35.06 | 9.50 | *p* < 0.001 |
| Cover Area (µm<sup>2</sup>) | 10840.47 | 3982.10 | 22043.11 | 7629.16 | *p* < 0.01 |
| Mean Bifurcation Angle (°) | 79.35 | 8.36 | 86.91 | 7.40 | *p* > 0.05 |
| Partition Asymmetry | 0.60 | 0.11 | 0.58 | 0.09 | *p* > 0.05 |
| Max Euclidean Distance (µm) | 41.41 | 10.76 | 75.16 | 22.69 | *p* < 0.01 |
| Max Path Distance (µm) | 47.53 | 10.83 | 91.44 | 23.97 | *p* < 0.001 |
| **(7 h)**<br>**Morphometric Feature** | **Carbon**<br>**Replete**<br>**(Mean)** | **±**<br>**Standard**<br>**Deviation** | **Carbon**<br>**Deplete**<br>**(Mean)** | **±**<br>**Standard**<br>**Deviation** | ***t*-test *p*-**<br>**value** |
| Thallus Diameter ($\mu\text{m}^2$) | 179.38 | 28.07 | 85.36 | 85.36 | $p < 0.001$ |
| Number of Bifurcations | 25.67 | 6.08 | 25.56 | 25.56 | $p > 0.05$ |
| Number of Tips | 28.11 | 6.33 | 26.89 | 26.89 | $p > 0.05$ |
| Width ($\mu\text{m}$) | 173.98 | 19.14 | 185.34 | 185.34 | $p > 0.05$ |
| Height ($\mu\text{m}$) | 166.68 | 26.11 | 215.32 | 215.32 | $p < 0.05$ |
| Depth ($\mu\text{m}$) | 10.88 | 3.14 | 10.85 | 10.85 | $p > 0.05$ |
| Total Length ($\mu\text{m}$) | 635.08 | 135.09 | 800.14 | 800.14 | $p < 0.05$ |
| Surface Area ($\mu\text{m}^2$) | 2394.43 | 484.77 | 2914.51 | 2914.51 | $p > 0.05$ |
| Volume ($\mu\text{m}^3$) | 982.62 | 190.12 | 946.72 | 946.72 | $p > 0.05$ |
| Rhizoidal Growth Unit ($\mu\text{m}$) | 23.00 | 4.75 | 31.23 | 31.23 | $p < 0.05$ |
| Cover Area ($\mu\text{m}^2$) | 29227.45 | 6934.82 | 39176.52 | 39176.52 | $p < 0.05$ |
| Mean Bifurcation Angle ($^\circ$) | 82.19 | 6.23 | 88.03 | 88.03 | $p < 0.05$ |
| Partition Asymmetry | 0.63 | 0.07 | 0.65 | 0.65 | $p > 0.05$ |
| Max Euclidean Distance ($\mu\text{m}$) | 62.15 | 7.66 | 120.92 | 120.92 | $p < 0.01$ |
| Max Path Distance ($\mu\text{m}$) | 75.05 | 5.18 | 156.50 | 156.50 | $p < 0.001$ |
| **(24 h)**<br>**Morphometric Feature** | **Carbon Replete (Mean)** | **± Standard Deviation** | **Carbon Deplete (Mean)** | **± Standard Deviation** | **t-test p-value** |
| Thallus Diameter ($\mu\text{m}^2$) | 2038.41 | 336.41 | 180.49 | 24.79 | $p < 0.01$ |
| Number of Bifurcations | 365.75 | 80.22 | 88.00 | 24.42 | $p < 0.01$ |
| Number of Tips | 433.25 | 106.58 | 90.63 | 23.74 | $p < 0.01$ |
| Width ($\mu\text{m}$) | 402.03 | 28.77 | 364.64 | 48.94 | $p < 0.05$ |
| Height ($\mu\text{m}$) | 401.90 | 15.90 | 393.74 | 19.71 | $p > 0.05$ |
| Depth ($\mu\text{m}$) | 25.14 | 5.19 | 9.69 | 2.75 | $p < 0.01$ |
| Total Length ($\mu\text{m}$) | 9918.81 | 2094.98 | 3015.64 | 815.10 | $p < 0.01$ |
| Surface Area ($\mu\text{m}^2$) | 43028.44 | 9579.73 | 12093.19 | 3594.02 | $p < 0.01$ |
| Volume ($\mu\text{m}^3$) | 28425.62 | 4653.46 | 4245.75 | 1349.18 | $p < 0.01$ |
| Rhizoidal Growth Unit ($\mu\text{m}$) | 23.16 | 1.92 | 33.23 | 2.53 | $p < 0.001$ |
| Cover Area ($\mu\text{m}^2$) | 161828.23 | 16638.78 | 143817.79 | 22032.07 | $p > 0.05$ |
| Mean Bifurcation Angle ($^\circ$) | 68.06 | 3.21 | 81.69 | 2.63 | $p < 0.01$ |
| Partition Asymmetry | 0.64 | 0.00 | 0.67 | 0.06 | $p > 0.05$ |
| Max Euclidean Distance ($\mu\text{m}$) | 206.91 | 18.27 | 181.53 | 34.94 | $p > 0.05$ |
| Max Path Distance ($\mu\text{m}$) | 256.90 | 24.26 | 235.21 | 82.52 | $p > 0.05$ |

**Supplementary Table 4.** Morphometric features of *R. globosum* rhizoids growing on chitin beads, associated with Figure 4 A-B.

| **Particulate Carbon**<br>**Morphometric Feature** | **(1 h)**<br>**(Mean)** | **±**<br>**Standard**<br>**Deviation** | **(4 h)**<br>**(Mean)** | **±**<br>**Standard**<br>**Deviation** | **(7 h)**<br>**(Mean)** | **±**<br>**Standard**<br>**Deviation** | **(24 h)**<br>**(Mean)** | **±**<br>**Standard**<br>**Deviation** |
| --- | --- | --- | --- | --- | --- | --- | --- | --- |
| Thallus Diameter ($\mu\text{m}^2$) | 71.38 | 12.81 | 83.29 | 9.78 | 91.89 | 16.18 | 269.71 | 55.76 |
| Number of Bifurcations | 2.22 | 1.30 | 8.89 | 4.70 | 10.75 | 4.83 | 112.75 | 62.47 |
| Number of Tips | 3.22 | 1.30 | 10.89 | 5.49 | 13.50 | 6.07 | 143.50 | 91.81 |
| Width ($\mu\text{m}$) | 131.18 | 14.57 | 152.85 | 30.43 | 153.14 | 17.21 | 171.85 | 15.96 |
| Height ($\mu\text{m}$) | 97.68 | 23.73 | 105.45 | 22.29 | 121.57 | 33.43 | 145.96 | 28.79 |
| Depth ($\mu\text{m}$) | 6.92 | 2.26 | 32.81 | 15.63 | 42.38 | 9.70 | 58.99 | 10.02 |
| Total Length ($\mu\text{m}$) | 32.95 | 16.05 | 191.55 | 73.31 | 324.11 | 108.77 | 2160.80 | 722.46 |
| Surface Area ($\mu\text{m}^2$) | 151.11 | 48.92 | 715.97 | 237.15 | 1164.11 | 457.21 | 7260.26 | 3195.89 |
| Volume ($\mu\text{m}^3$) | 104.61 | 26.55 | 291.69 | 96.99 | 422.49 | 177.54 | 2740.66 | 1364.26 |
| Rhizoidal Growth Unit ($\mu\text{m}$) | 10.50 | 4.15 | 18.97 | 5.76 | 25.68 | 6.64 | 17.54 | 5.72 |
| Cover Area ($\mu\text{m}^2$) | 12718.87 | 2934.84 | 15959.67 | 4130.87 | 18483.84 | 4996.53 | 24937.91 | 4319.79 |
| Mean Bifurcation Angle ($^\circ$) | 94.20 | 18.51 | 81.76 | 12.22 | 92.12 | 9.37 | 82.45 | 1.73 |
| Partition Asymmetry | 0.37 | 0.28 | 0.40 | 0.19 | 0.56 | 0.07 | 0.58 | 0.12 |
| Max Euclidean Distance ($\mu\text{m}$) | 10.43 | 5.91 | 52.95 | 31.98 | 43.05 | 17.77 | 41.48 | 15.56 |
| Max Path Distance ($\mu\text{m}$) | 12.63 | 7.17 | 88.61 | 61.80 | 66.92 | 21.46 | 63.91 | 15.28 |

**Supplementary Table 5.** Morphometric features and statistical comparisons of searching *R. globosum* rhizoids encountering chitin beads, associated with Figure 4 D-E.

| **Rhizoid Differentiation**<br>**Morphometric Feature** | **Particle**<br>**Associated**<br>**(Mean)** | **±**<br>**Standard**<br>**Deviation** | **Not Particle**<br>**Associated**<br>**(Mean)** | **±**<br>**Standard**<br>**Deviation** | **t-test *p*-**<br>**value** |
| --- | --- | --- | --- | --- | --- |
| Number of Bifurcations | 24.63 | 11.33 | 28.50 | 8.65 | $p > 0.05$ |
| Number of Tips | 30.50 | 12.75 | 30.25 | 8.43 | $p > 0.05$ |
| Width (µm) | 150.84 | 22.60 | 217.94 | 10.43 | $p < 0.001$ |
| Height (µm) | 146.25 | 31.11 | 201.50 | 29.49 | $p < 0.001$ |
| Depth (µm) | 33.32 | 6.52 | 9.49 | 5.17 | $p < 0.001$ |
| Total Length (µm) | 465.18 | 167.59 | 1090.68 | 310.27 | $p < 0.01$ |
| Surface Area (µm<sup>2</sup>) | 1415.67 | 623.56 | 3913.74 | 1420.74 | $p < 0.01$ |
| Volume (µm<sup>3</sup>) | 406.93 | 236.86 | 1291.92 | 595.80 | $p < 0.001$ |
| Rhizoidal Growth Unit (µm) | 15.88 | 4.42 | 36.16 | 4.44 | $p < 0.001$ |
| Cover Area (µm<sup>2</sup>) | 22093.22 | 6323.01 | 43929.59 | 6729.19 | $p < 0.001$ |
| Mean Bifurcation Angle (°) | 85.71 | 9.99 | 92.76 | 7.28 | $p < 0.001$ |
| Partition Asymmetry | 0.54 | 0.11 | 0.53 | 0.15 | $p > 0.05$ |
| Max Euclidean Distance (µm) | 32.80 | 11.82 | 127.76 | 27.94 | $p < 0.001$ |
| Max Path Distance (µm) | 85.76 | 41.84 | 171.28 | 57.36 | $p < 0.05$ |

**Supplementary Movie 1 – 4D imaging of developing *R. globosum* rhizoids used for quantifying morphometric growth trajectories (Replicate 1).** Time in HH:MM

**Supplementary Movie 2 – 4D imaging of developing *R. globosum* rhizoids used for quantifying morphometric growth trajectories (Replicate 2).** Time in HH:MM

**Supplementary Movie 3 – 4D imaging of developing *R. globosum* rhizoids used for quantifying morphometric growth trajectories (Replicate 3).** Time in HH:MM

**Supplementary Movie 4 – 4D imaging of developing *R. globosum* rhizoids used for quantifying morphometric growth trajectories (Replicate 4).** Time in HH:MM

**Supplementary Movie 5 – 4D imaging of developing *R. globosum* rhizoids used for quantifying morphometric growth trajectories (Replicate 5).** Time in HH:MM

**Supplementary Movie 6 – Representative 3D reconstructions of 7 h *R. globosum* rhizoids from caspofungin treated and control cells.** Cell wall inhibited rhizoids display atypical hyperbranching.

**Supplementary Movie 7 – Representative 3D reconstructions of 7 h *R. globosum* rhizoids from cytochalasin B treated and control cells.** Actin inhibited rhizoids display atypical hyperbranching.

**Supplementary Movie 8 – 4D imaging of the entire *R. globosum* life cycle growing on 10 mM NAG.** Cell completes its entire lifecycle and sporulates. Time in HH:MM

**Supplementary Movie 9 – 4D imaging of *R. globosum* growing in carbon deplete media.** Cell does not complete lifecycle and ceases growth after 14-16 h. Time in HH:MM

**Supplementary Movie 10 – Representative 3D reconstructions of *R. globosum* rhizoids from carbon replete and carbon deplete cells.** Cells in the carbon deplete condition display the differential searching phenotype. Reconstructions are scaled relative to timepoint.

**Supplementary Movie 11 – Representative 3D reconstructions of *R. globosum* rhizoids from cells growing on chitin beads.** Reconstructions are scaled relative to timepoint.

**Supplementary Movie 12 – 4D imaging of *R. globosum* growing on a chitin microbead.** Note that branching within the bead emanates from ‘pioneer’ penetrative rhizoids. Time in HH:MM

**Supplementary Movie 13 – 4D imaging of searching *R. globosum* rhizoids encountering a chitin bead (XY).** Note how rhizoids not in contact with the particle continue to grow in a searching pattern. Time in HH:MM

**Supplementary Movie 14 – 4D imaging of searching *R. globosum* rhizoids encountering a chitin bead (YZ).** Note how branching is most profuse in rhizoids in contact with the particle. Time in HH:MM

**Supplementary Movie 15 – Representative 3D reconstruction of *R. globosum* rhizoids from a searching cell in carbon deplete media that has encountered a chitin microbead.** The rhizoid is spatially differentiated and coloured whether in contact (green) or not in contact with (blue) the chitin bead.

Supplementary File 1 – Total raw data used for analysis in this study

## Notes

#### Summary of Updates

Supplementary Figure 7 corrected

